# Probabilistic uncertainty in virtual spatial transcriptomics from histology is largely redundant with the predicted mean

**DOI:** 10.64898/2026.10.23.760875

**Authors:** Thomas Batique, Kalin Nonchev, Viktor H Koelzer, Gunnar Rätsch

## Abstract

Deep learning has enabled virtual spatial transcriptomics from histology, offering a scalable approach to molecular profiling of routinely archived tissue without molecular assays. However, reliable uncertainty estimates are essential for identifying potentially inaccurate predictions, and it remains unclear whether existing methods provide useful information about prediction reliability. Here, we present a systematic benchmark of uncertainty estimation for virtual spatial transcriptomics across three cancer cohorts spanning melanoma, lung, and kidney tumors. We evaluate aleatoric and epistemic uncertainty using probabilistic likelihoods and approximate Bayesian methods alongside histology-free and training-free baselines, assessing both calibration and error informativeness. Across cohorts and methods, uncertainty is associated with prediction error, but the predicted mean largely explains this association. After accounting for the mean, uncertainty provides only weak information about larger errors. Moreover, good calibration does not imply informative uncertainty, with simple distributions fitted without histology achieving comparable calibration. We therefore distinguish calibrated from informative uncertainty and argue that uncertainty should provide information about prediction error beyond the predicted mean. This benchmark provides a framework for systematic uncertainty evaluation in virtual spatial transcriptomics.

**Code availability:** https://github.com/ratschlab/uncertainty-vst-2026

## 1 Introduction

Cancer tissue is routinely archived as hematoxylin and eosin (H&E)-stained histology, yet spatial molecular measurements are unavailable for most specimens. Spatial transcriptomics (ST) measures gene expression while preserving its spatial context [1], but current technologies remain costly and require specialized tissue processing. Recently, advances in deep learning have enabled the prediction of spatial gene expression directly from histology [2–6]. Related approaches have further increased the spatial resolution of these predictions by inferring expression at subspot or single-cell resolution from histology [7, 8]. If reliable, virtual spatial transcriptomics could enable molecular characterization of routinely archived clinical tissue, supporting large-scale studies of disease biology.

Typically, these models return a single expression estimate for each spot and gene, without quantifying predictive uncertainty [9]. This makes it difficult to assess the reliability of individual predictions and limits the use of virtual spatial transcriptomics in applications such as molecular subtyping, biomarker discovery, and characterization of spatially heterogeneous tumor regions. Per-prediction uncertainty estimates could provide a measure of confidence and help distinguish predictions that are likely to be accurate from those that are less reliable.

Alternatively, probabilistic models can provide such estimates by predicting a distribution rather than a single value, from which predictive uncertainty can be quantified. For example, in medical imaging classification, uncertainty-based deferral can improve accuracy [10], including for H&E slides [11]. Similar benefits have been reported for medical regression [12]. However, uncertainty does not consistently identify prediction errors across domains, with estimates in molecular property regression failing to rank prediction errors reliably across datasets [13]. In spatial transcriptomics, several H&E-based models predict uncertainty alongside the expression mean, but whether these estimates provide useful information about prediction accuracy remains largely unexplored [3, 14, 15].

A key challenge arises from the distribution of gene expression itself. Specifically, count data are overdispersed, with variance that typically increases with the mean [16, 17], a relationship further assumed in models of expression [18, 19]. Predicted variance may therefore depend strongly on the predicted mean. Consequently, if prediction error also follows the mean, an association between uncertainty and error may reflect that shared dependence rather than information about which individual predictions are less reliable. Therefore, we distinguish between two properties of predictive uncertainty: (1) calibration, which examines whether predicted distributions accurately represent observed outcomes, and (2) informativeness, which assesses whether uncertainty identifies predictions associated with larger errors.

To address this, we introduce a systematic benchmark for predictive uncertainty in virtual spatial transcriptomics from histology. The benchmark spans probabilistic likelihoods, epistemic uncertainty estimators, and histology-free and training-free baselines, evaluated on three Visium cohorts covering melanoma, lung, and kidney tumors. Specifically, it evaluates both calibration and the informativeness of uncertainty for identifying prediction errors beyond those explained by the predicted mean. To our knowledge, this is the first systematic evaluation of predictive uncertainty in virtual spatial transcriptomics.

Across cohorts and methods, we find that uncertainty is associated with prediction error, but the predicted mean largely explains this association. After accounting for the mean, uncertainty identifies larger errors only marginally. Moreover, distributions fitted without histology achieve comparable calibration, showing that good calibration does not necessarily imply informative uncertainty. These results highlight a distinction between calibrated uncertainty, which captures expected expression variability, and informative uncertainty, which provides information about individual prediction errors.

## 2 Related work

### 2.1 Virtual spatial transcriptomics from H&E images

Deep learning models have been developed to infer spatial gene expression from H&E images using convolutional, graph-based, transformer, and generative architectures, as well as pathology foundation model embeddings [2–6]. These models exploit morphological features to predict spatial patterns of gene expression, with prediction accuracy varying substantially across genes and tissue contexts [4]. Beyond expression prediction, related work explores the link between morphology and molecular state through generative models that synthesize histopathology images conditioned on morphology embeddings [20] or transcriptomic profiles [21], and language-based systems that make molecular information queryable from histopathology images [22, 23]. However, like existing expression predictors, these approaches do not quantify uncertainty for individual predictions. Current evaluations primarily focus on the accuracy of the predicted mean, typically using per-gene correlation with measured expression. Here, we instead focus on the uncertainty associated with these predictions.

### 2.2 Probabilistic uncertainty in deep learning

A regression network trained with squared error loss produces a point prediction, whereas a probabilistic readout predicts the parameters of an output distribution, typically by maximum likelihood. Predictive uncertainty is commonly decomposed into aleatoric and epistemic components [24]. Aleatoric uncertainty captures variability in the data-generating process and is represented by the spread of the predictive distribution. Epistemic uncertainty reflects uncertainty in the model and can decrease as the amount of training data increases. Common approximations include deep ensembles [25], Monte Carlo dropout [26], and Laplace approximations [27].

Calibration assesses whether predicted distributions agree with observed outcomes in aggregate [28], but it does not imply that uncertainty is informative about individual prediction errors. For example, a model that ignores its input and predicts the marginal distribution of the target can achieve good calibration without indicating which individual predictions are likely to be inaccurate [29]. Calibration alone therefore does not establish that uncertainty can identify unreliable predictions.

### 2.3 Uncertainty in virtual spatial transcriptomics

Existing approaches to uncertainty in virtual spatial transcriptomics fall into two settings. The first concerns predictive uncertainty estimated directly by the model. A few H&E-based methods predict a variance alongside the expression mean, but whether these estimates are informative about prediction error has not been systematically evaluated. For example, STimage reports that its predicted variance increases with the predicted mean [14]. Hist2ST predicts distributional parameters but does not evaluate whether they provide useful information about prediction error [3]. A recent Bayesian neural network study reports enrichment of prediction errors at both ends of the uncertainty ranking, but does not account for the dependence between uncertainty and expression level [15].

The second setting concerns post hoc uncertainty estimates constructed from model predictions. Conformal prediction can be used to construct prediction intervals with distribution-free coverage guarantees. For example, TISSUE derives an initial uncertainty estimate from variability among a cell’s nearest spatial neighbors and uses it to construct prediction intervals calibrated against measured genes [30]. However, TISSUE targets transcripts imputed from a paired single-cell reference rather than expression predicted directly from histology. UTOPIA operates on virtual spatial transcriptomics from H&E images [31], but assigns confidence to a detection decision under false discovery rate control rather than to a continuous expression prediction.

Here, we focus on model-based predictive uncertainty, where the model estimates a predictive distribution and its associated uncertainty directly from the input.

## 3 Methods

### 3.1 Task, data, and model

We evaluate predictive uncertainty for virtual spatial transcriptomics by predicting a distribution for each spot-gene pair (fig. 1). Given an H&E image patch centered at spot *s*, the model predicts a distribution over the expression *y*_*sg*_ of gene *g*.

**Figure 1:**
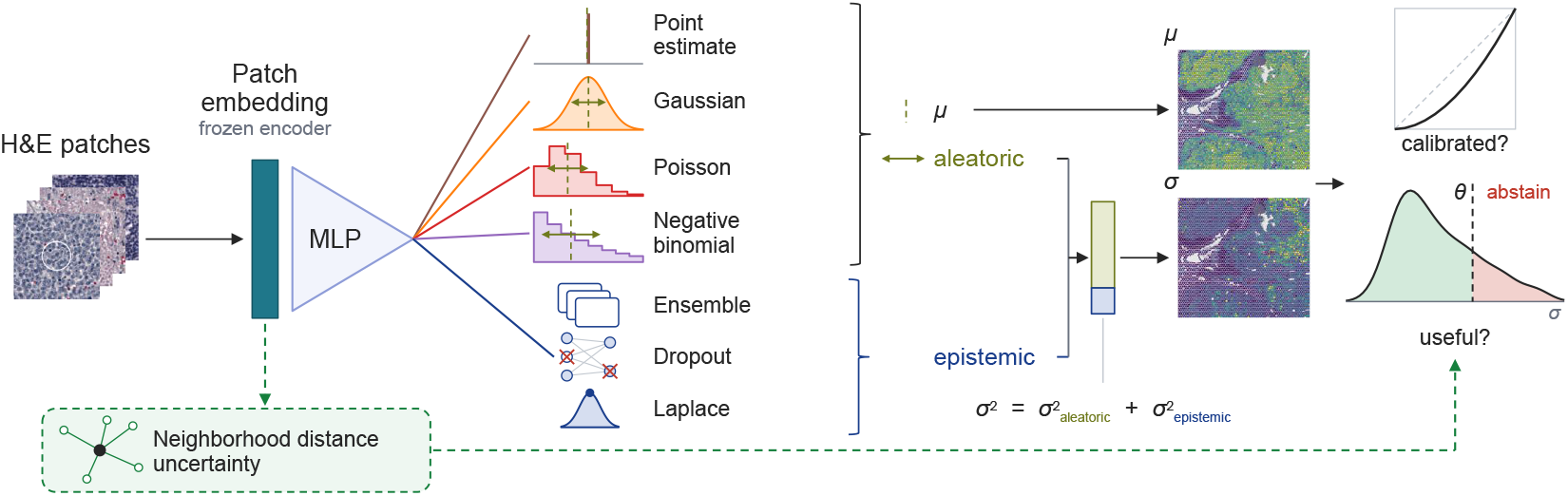
Graphical abstract. An MLP maps frozen H&E patch embeddings to the parameters of a predictive distribution. The aleatoric spread comes from the head, the epistemic spread from three epistemic estimators. A training-free neighborhood signal bypasses the model. We assess calibration and informativeness.

We use three Visium cohorts: Tumor Profiler (18 slides, 7 donors, metastatic melanoma), Kidney/Lung TLS (6 slides, 6 donors, kidney and lung tumors enriched for tertiary lymphoid structures), and NSCLC (36 slides, 8 donors, non-small cell lung carcinoma). Each cohort is restricted to its 3,000 most highly variable genes. Evaluation uses leave-one-patient-out cross-validation, with each donor held out once. Scores are summarized across donors using a Student’s *t* 95% confidence interval. Cohort descriptions and preprocessing are provided in section B.

Each spot is represented by a frozen Midnight-12k embedding [32] of an H&E patch spanning 1.5 spot diameters, following ST-Net [2]. A multilayer perceptron maps the embedding to gene-specific distribution parameters and is trained by minimizing the negative log-likelihood. The model trunk is fixed after a single architecture search, so all models use the same image representation and prediction network and differ only in their probabilistic readout and uncertainty estimation.

### 3.2 Probabilistic readouts and sources of uncertainty

We evaluate three likelihood families that differ in how aleatoric variance depends on the predictive mean,

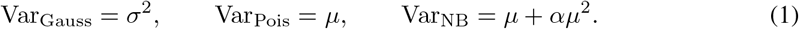

The Gaussian variance is independent of the mean, whereas the Poisson and negative binomial variances are mean-dependent. The negative binomial also has a dispersion parameter *α*.

We also evaluate two constrained Gaussian models, with variance held constant across spots within each gene or across genes within each spot. The three Gaussian variants are referred to as *heteroscedastic, gene-homoscedastic*, and *spot-homoscedastic*. Gaussian models are trained on a depth-corrected log target, while count-based readouts are trained on integer counts normalized to a common sequencing depth. All mean predictions are evaluated on the common scale log(1 + counts per 10^4^) (section D).

Aleatoric uncertainty is the variance of the predictive distribution at fixed model parameters, while epistemic uncertainty, estimated using deep ensembles, Monte Carlo dropout, and a last-layer Laplace approximation, captures uncertainty in the model itself. The two combine through the law of total variance to obtain total predictive uncertainty, with the construction of every estimator given in section D.

### 3.3 Evaluation

#### Accuracy

Pearson correlation between prediction and measurement is computed per gene within each slide, averaged over slides within each donor and then over genes. This slide-aware Pearson correlation avoids between-slide mean shifts and prevents donors with multiple slides from being overrepresented. Slides with constant predictions or targets are excluded.

#### Calibration

Calibration is assessed using the probability integral transform, in the nonrandomized form of Czado et al. [33], which works for continuous and discrete predictives alike. Since the transform is invariant to strictly increasing reparameterizations, the Gaussian and count heads are compared under one diagnostic despite their different targets. Calibration error is the mean absolute deviation between empirical and nominal coverage across probability levels.

#### Association, informativeness, and selective utility

Writing *u* for predictive standard deviation and Δ for absolute error, we measure the association between uncertainty and error using Spearman correlation in two groupings. The first ranks errors across spots within each gene and slide, and the second ranks errors across genes within each spot and slide. Within a group, we report the unconditional correlation *ρ*_*u*Δ_ and the partial Spearman correlation controlling for the predicted mean μ,

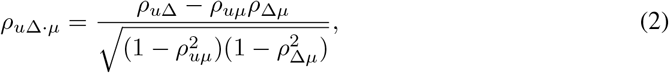

where all correlations are computed on the same entries and averaged within each held-out donor. The partial correlation tests whether uncertainty provides information about prediction error beyond the predicted mean.

Selective utility is assessed operationally, by withholding the least certain predictions and recomputing the error on the remaining predictions. Results are compared with an oracle that withholds predictions according to their true absolute error. Section E gives the grouping rules and withholding protocol.

### 3.4 Baselines

The *mean-only* baseline is trained with squared error loss using the same model trunk and target. The *marginal* baseline uses no histology input: it fits one expression distribution per gene on the training patients and assigns it to every spot in the held-out patient. Because it produces no spot-specific prediction, its slide-aware Pearson correlation is undefined, so it provides a calibration reference instead.

The *neighborhood* baseline is training-free: it uses the dispersion of measured expression among a spot’s nearest neighbors in the frozen embedding space as an aleatoric signal and the distance to those neighbors as an epistemic signal [13]. The *oracle* ranks predictions by their true absolute error and provides an upper bound for uncertainty-based ranking. Finally, *STimage* [14] is reproduced as a linear head trained as a deep ensemble using the same embeddings, splits, and scoring procedures.

## 4 Experiments

### 4.1 Probabilistic modeling preserves prediction accuracy

Probabilistic readouts achieve similar prediction accuracy to the mean-only baseline, indicating that modeling predictive variance adds little cost to the mean prediction, with the Poisson model on NSCLC as the only exception (section A.1). Overall predictive performance is modest across cohorts (fig. 4), with the best-performing probabilistic readouts among the 50 most predictive genes achieving slide-aware Pearson correlations of only 0.55, 0.26, and 0.24 on Tumor Profiler, Kidney/Lung TLS, and NSCLC, respectively. Prediction accuracy is strongly associated with mean expression, with highly expressed genes generally being better predicted (section A.1).

### 4.2 Calibration depends primarily on the likelihood family

Figure 2 shows calibration error for each likelihood family, computed from the aleatoric uncertainty alone and averaged over the *n* most predictive genes. Epistemic uncertainty is substantially smaller than aleatoric uncertainty and has little effect on calibration. Across all 3,000 genes, the ranking is consistent across cohorts, with the negative binomial best calibrated, followed by the Poisson and Gaussian variants.

**Figure 2:**
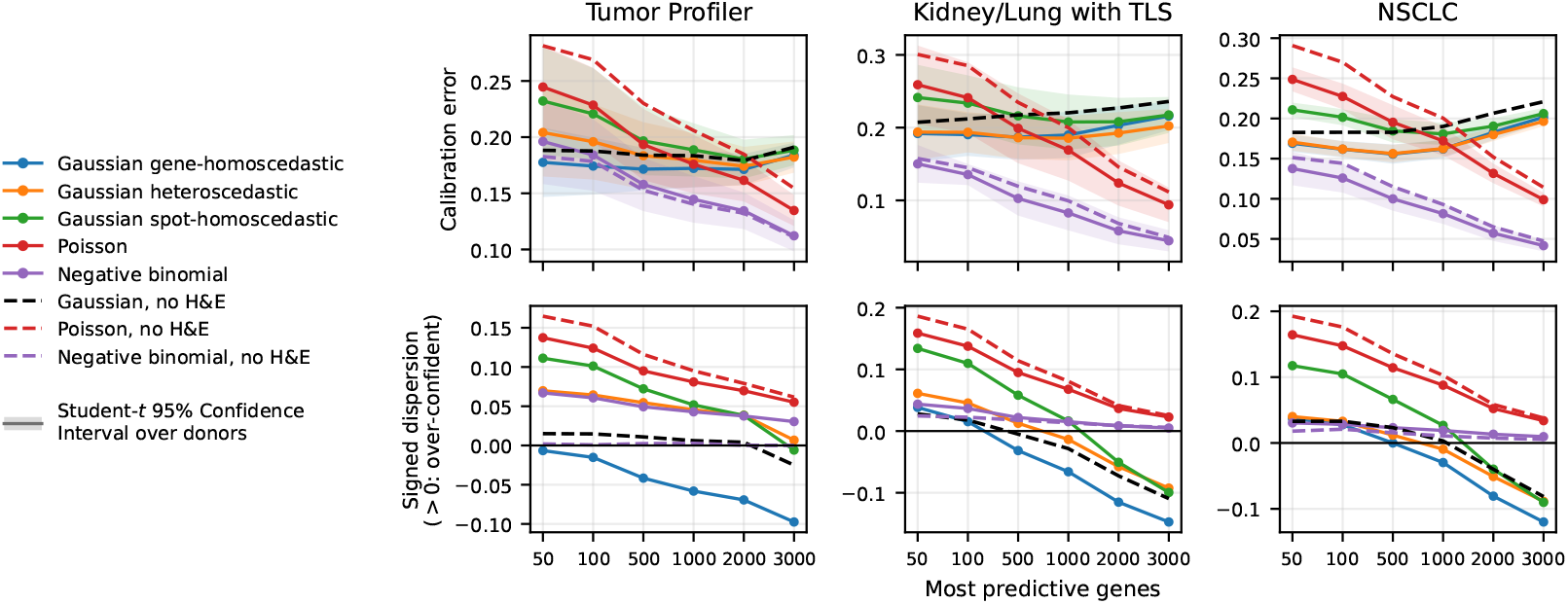
Calibration of each likelihood family (solid lines) against a marginal baseline (dashed lines) of the same family fitted to measured expression without using histology. Columns show the three cohorts. Each metric is averaged over the *n* most predictive genes according to a shared ranking. *Top row:* calibration error. *Bottom row:* signed dispersion, with positive values indicating an overly narrow predictive distribution.

Among the 50 most predictive genes, the ranking changes. The negative binomial has calibration error comparable to the heteroscedastic and gene-homoscedastic Gaussian models, while the spot-homoscedastic Gaussian and Poisson models are less well calibrated. Gaussian calibration remains relatively stable across genes, whereas calibration of the count likelihoods improves for genes with lower expression, which are also less accurately predicted. The Gaussian models are overconfident for the best-predicted genes and underconfident across the full gene panel (bottom row, fig. 2), so a single multiplicative adjustment to the predicted variance cannot correct both regimes.

The marginal baseline reproduces the same ranking of likelihood families (dashed lines, fig. 2), indicating that similar calibration can be obtained without using histology. Relative to the corresponding marginal baseline, the Poisson shows the largest improvement across cohorts, the Gaussian models show improvement in two cohorts, and the negative binomial shows no measurable improvement. In particular, the negative binomial marginal baseline, a simple baseline with no histology input at all, achieves low calibration error while assigning the same distribution to every spot. Thus, good calibration does not imply that uncertainty identifies individual prediction errors [29].

### 4.3 Uncertainty-error associations largely reflect the predicted mean

Figure 3 shows the Spearman correlation between predicted spread and absolute error before and after controlling for the predicted mean. Correlations are computed across the spots of each gene and across the genes of each spot.

**Figure 3:**
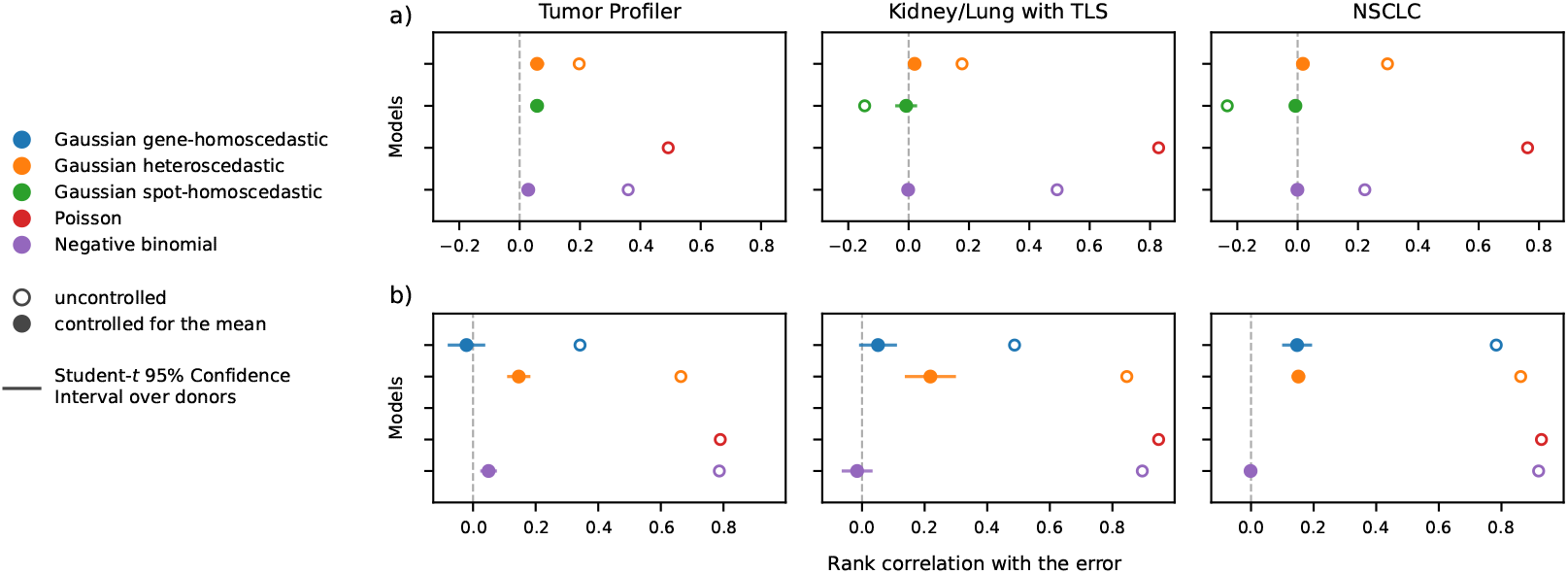
Spearman correlation between predicted uncertainty and absolute error, uncontrolled and using partial Spearman correlation controlling for the predicted mean. Correlations are computed across the spots of each gene (top row) and across the genes of each spot (bottom row). The Poisson has no controlled value because its variance is equal to its mean.

Across the genes of a spot, the uncontrolled correlation reaches 0.8 in all three cohorts, which, taken alone, would read as a strong reliability signal. Most of that association is explained by the predicted mean, the negative binomial correlation on Tumor Profiler decreasing from +0.79 to +0.05 once the mean is controlled for, and the other two cohorts behaving the same way.

This behavior is expected for the Poisson and partly expected for the negative binomial (eq. (1)). The heteroscedastic Gaussian shows the same behavior despite having a free variance. On Tumor Profiler, its correlation decreases from +0.66 to +0.15 after controlling for the mean. Correlations across the spots of a gene are smaller, but the same pattern persists. After controlling for the predicted mean, all correlations in this analysis lie between − 0.07 and +0.22. The residual association is strongest among the genes with the largest prediction errors (section A.5).

The other uncertainty measures do not provide a stronger signal (section A.4). The three epistemic estimators are mostly weaker than the aleatoric uncertainty, at best comparable to it, and inconsistent in direction across methods. The training-free neighborhood dispersion, computed from measured expression without fitting an uncertainty model, has correlations of only +0.01 to +0.06 across the genes of a spot. Neighborhood distance provides no measurable signal, staying below +0.025. The STimage readout shows the same pattern as our own heads, on both the depth-corrected and the raw count target (fig. 7). Since the two readouts share embeddings, likelihood, target, and splits, the result does not appear to depend on the readout architecture.

Figures 9 and 10 test whether withholding the least certain predictions improves the error of the remainder (sections A.6 and E). The oracles reduce the area under the curve by a third to a half, so prediction errors can in principle be ranked. On all entries, withholding within a (gene, slide) group, the predicted mean alone recovers as much of that reduction as the predicted uncertainty, or more (table 4). On the detected entries at a fixed predicted mean, the uncertainty recovers at most 30% of the oracle reduction. In absolute terms, withholding half of the predictions then lowers the mean error by at most 16%. The epistemic terms recover at most 7% of the oracle, and STimage behaves like our own heads.

### 4.4 The main result is robust to modeling choices

Selecting models on validation negative log-likelihood rather than on the Pearson correlation of the predicted mean reduces mean accuracy without improving informativeness, and the main finding is unchanged under the alternative *β*-NLL loss for the Gaussian heads [34] (section A.3).

Informativeness is unchanged under an alternative relative error definition (section E) and when the analysis is restricted to the *detected* entries, the spot-gene pairs whose measured count is non-zero (section A.3). That restriction is informative, because the uncontrolled correlations then become comparable to the controlled ones. The large uncontrolled correlations in the full data are therefore driven in substantial part by the zero entries.

Alternative uncertainty measures, including the coefficient of variation (predicted spread divided by predicted mean) and 1 + *α* for the negative binomial, also do not improve informativeness (section A.3).

Rounding the depth-corrected targets to integers, which the count likelihoods require, has no measurable effect on accuracy at the sequencing depths considered here (section B). Training the count heads on raw counts likewise does not improve the relationship between uncertainty and prediction error (section A.3).

Post hoc recalibration of the predicted variance does not improve informativeness either. Rescaling the variance per gene in bins of the predicted mean slightly lowers calibration error but does not improve any uncertainty-error correlation (section F).

### 4.5 Predicted uncertainty follows the mean-variance structure of the data

The measurements themselves show a strong mean-variance relationship. Genes with higher mean expression also exhibit higher variance across the spots of a slide, with a Spearman correlation of at least 0.95. Averaged across the spots of a slide, the predicted variance reproduces this ordering almost exactly, regardless of the prediction head.

This shared dependence on the predicted mean accounts for much of the uncontrolled uncertainty-error association in fig. 3. On all entries, across the genes of a spot, the heteroscedastic Gaussian’s predicted spread follows the predicted mean at 0.90 to 0.95, while absolute error follows the predicted mean at 0.67 to 0.89. Their product, 0.62 to 0.84, is comparable to the observed uncontrolled correlations of 0.66 to 0.86. Controlling for the predicted mean removes this shared association, leaving little residual correlation.

A complementary variance decomposition on the ranks splits a slide’s spot-by-gene matrix into gene, spot, and remainder terms. Only the remainder term varies from one prediction to the next. It accounts for 0.10 to 0.15 of the variance in predicted spread, against 0.40 to 0.70 of the variance in absolute error. Most of the variation in prediction error is not reflected in the remaining variation of uncertainty.

The two grouping schemes expose different aspects of this limitation. Across genes within a spot, predicted spread varies substantially but is strongly associated with the predicted mean. Across spots within a gene, that association is much weaker, but the remaining variation is small relative to the variation in error. From the tenth to the ninetieth percentile, predicted spread changes by a factor of only 1.40 to 1.73, whereas absolute error changes by a factor of 4.5 to 11.4.

#### Boundary conditions

The finding extends across every uncertainty measure evaluated. The fitted variance is largely redundant with the predicted mean, while the epistemic terms and the neighborhood distance provide little measurable error signal even before controlling for the mean. It is not limited to count targets, since the heteroscedastic Gaussian behaves the same way on a log-transformed target with a free variance. Our post hoc recalibration does not alter this pattern (section F).

The predicted spread remains informative in some settings. Although it does not reliably distinguish individual prediction errors, it retains information about tissue type at matched predicted expression levels (section A.7). Under image artifacts, the ensemble and distance-based uncertainty measures also increase and can flag degraded inputs (section A.7), suggesting that uncertainty may capture image-dependent variation under sufficiently distinct conditions, even when it provides little information about typical prediction errors.

Uncertainty should therefore be assessed on informativeness, its association with error after controlling for the predicted mean, and on selective utility, through sparsification against an oracle. Calibration and the uncontrolled uncertainty-error correlation alone can both be achieved by an uncertainty estimate that primarily reflects the predicted mean.

## 5 Discussion and conclusion

We presented a systematic benchmark of predictive uncertainty for virtual spatial transcriptomics and evaluated whether uncertainty identifies unreliable expression predictions. Across three cohorts, probabilistic readouts preserve prediction accuracy and produce generally well-calibrated predictive distributions. However, uncertainty is strongly associated with the predicted mean, and this association largely explains its relationship with absolute prediction error. After controlling for the predicted mean, uncertainty identifies predictions with larger errors only marginally. This pattern is consistent across likelihoods and epistemic uncertainty estimators, and extends to previously proposed and training-free uncertainty approaches evaluated in the same framework.

Specifically, the observed mean-variance relationship is already present in the measured expression data before model fitting. This is consistent with the fitted variance capturing expression-dependent variability rather than image-dependent prediction error. It also explains how a predictive distribution can be well calibrated while providing little information about the reliability of individual predictions.

These findings differ from settings such as medical imaging classification, where uncertainty can support selective prediction by identifying cases on which a model is likely to be wrong [11]. In regression, uncertainty that primarily follows the predicted mean may instead reflect the expected variability of the target rather than prediction reliability. Our results therefore highlight an important distinction between calibrated and informative uncertainty. For virtual spatial transcriptomics, uncertainty should be evaluated not only by calibration, but also by whether it identifies predictions with larger errors beyond what is explained by the predicted mean [29]. This benchmark provides a framework for making that distinction and motivates future uncertainty models that capture image-dependent sources of prediction error.

## 6 Limitations and future directions

Several limitations constrain our conclusions. First, evaluation is primarily in distribution, using held-out patients from the same cohorts, with no real batch, scanner, or staining shifts. Synthetic image artifacts (section A.7) offer a partial proxy, showing that distance-based and ensemble uncertainty respond to distribution shift even though they do not rank in-distribution errors. Established protocols for evaluating robustness to H&E staining variation [35] would provide a more realistic test of whether uncertainty tracks such shifts. Second, the measured ST expression is itself subject to technical and biological variability [36–38], such that observed prediction error does not exclusively reflect model error. Without replicate measurements, these sources of measurement variability cannot be disentangled from model error. Third, we use a single frozen image encoder, so epistemic uncertainty captures the expression readout but not representation uncertainty. Finally, the model trunk is tuned once using a mean-only objective and then fixed, so our models are not optimized for state-of-the-art expression prediction (section C).

Our main conclusion concerns error informativeness rather than the magnitude of uncertainty-error correlations. Sparsification recovers only a small fraction of the improvement achievable using true errors.

Future work should develop uncertainty estimates that capture image-dependent prediction error rather than mean-dependent variation. Distribution-free approaches [39], including conformal methods in spatial transcriptomics [30, 31], provide one possible direction. More targeted recalibration, conditioned on covariates rather than on the predicted mean alone, provides another [40, 41], although the recalibrations we test do not improve error informativeness (section F).

## Acknowledgments and Disclosure of Funding

Computational data analysis was performed at Leonhard Med (https://sis.id.ethz.ch/services/sensitiveresearchdata/), a secure trusted research environment at ETH Zurich. We thank the patients and their families and the clinical and laboratory teams supporting them, whose contributions made this study possible. We also use data from the Tumor Profiler (TuPro) study [42] and thank the Tumor Profiler Consortium (https://tumorprofilercenter.ch).

## Ethics statement

This work reanalyzes histology and spatial transcriptomics data from cohorts collected and consented under the ethical approvals of their original studies [43–45]. No new human tissue or patient data were collected for this work. Kidney/Lung TLS and NSCLC are publicly available; Tumor Profiler is accessible upon request at https://tumorprofilercenter.ch/contacts. Virtual spatial transcriptomics could eventually inform clinical decisions, and our central finding argues for caution in that setting. Predicted uncertainty in this task does not reliably identify individual unreliable predictions, so per-prediction confidence from these models should not be used to guide clinical decisions without further validation.

## Reproducibility statement

Code implementing the models, training procedures, and evaluation metrics is available at the link given in the abstract. Cohort composition, preprocessing, and target construction are described in section B. Model architecture, training procedure, hyperparameter search, and compute are described in section C. The construction of every uncertainty estimator is given in section D, and every evaluation metric, including calibration, error informativeness, and the grouping and withholding rules for sparsification, is given in section E. Ensemble members differ only in random seed, and the hyperparameter search was run under a fixed seed (section C).

## Use of large language models

Generative AI tools (including large language models such as OpenAI ChatGPT and Anthropic Claude, as well as code-completion assistants such as GitHub Copilot) were used during the preparation of this manuscript to assist with prose editing, literature synthesis, and software development. All AI-generated content was critically reviewed, edited, and verified by the authors, who take full responsibility for the accuracy, integrity, and originality of the final work. No AI tool was used to generate scientific ideas, interpret results, or draw conclusions.

## A Additional analyses

### A.1 The cost of an uncertainty head on mean accuracy

Figure 4 compares each head to the mean-only baseline within each donor. The cost in mean performance is small. On Tumor Profiler, on the full 3,000 gene panel, the maximum cost in accuracy relative to the mean-only baseline is 4%, reached on the Poisson and the negative binomial. On the same cohort, the Gaussians slightly outperform the mean-only baseline, by at most 2%. On NSCLC, on the 3,000 gene panel, the cost in mean performance is at most 2%, except for the Poisson, which incurs a 27% cost. On the 50 best-predicted genes, the cost is at most 7%, reached for the negative binomial and the Poisson, and 4% for the heteroscedastic head. Finally, on Kidney/Lung TLS, a cohort with low overall mean accuracy, the cost in mean performance on the 3,000 gene panel is at most 15%, for the Poisson. We attribute these higher relative costs to that cohort’s lower overall mean accuracy. The Poisson must fit both the mean and the variance with a single parameter, which explains its degraded performance. Panel *c)* shows the mean expression along the same gene ordering. The genes with the highest mean are also those best predicted.

**Figure 4:**
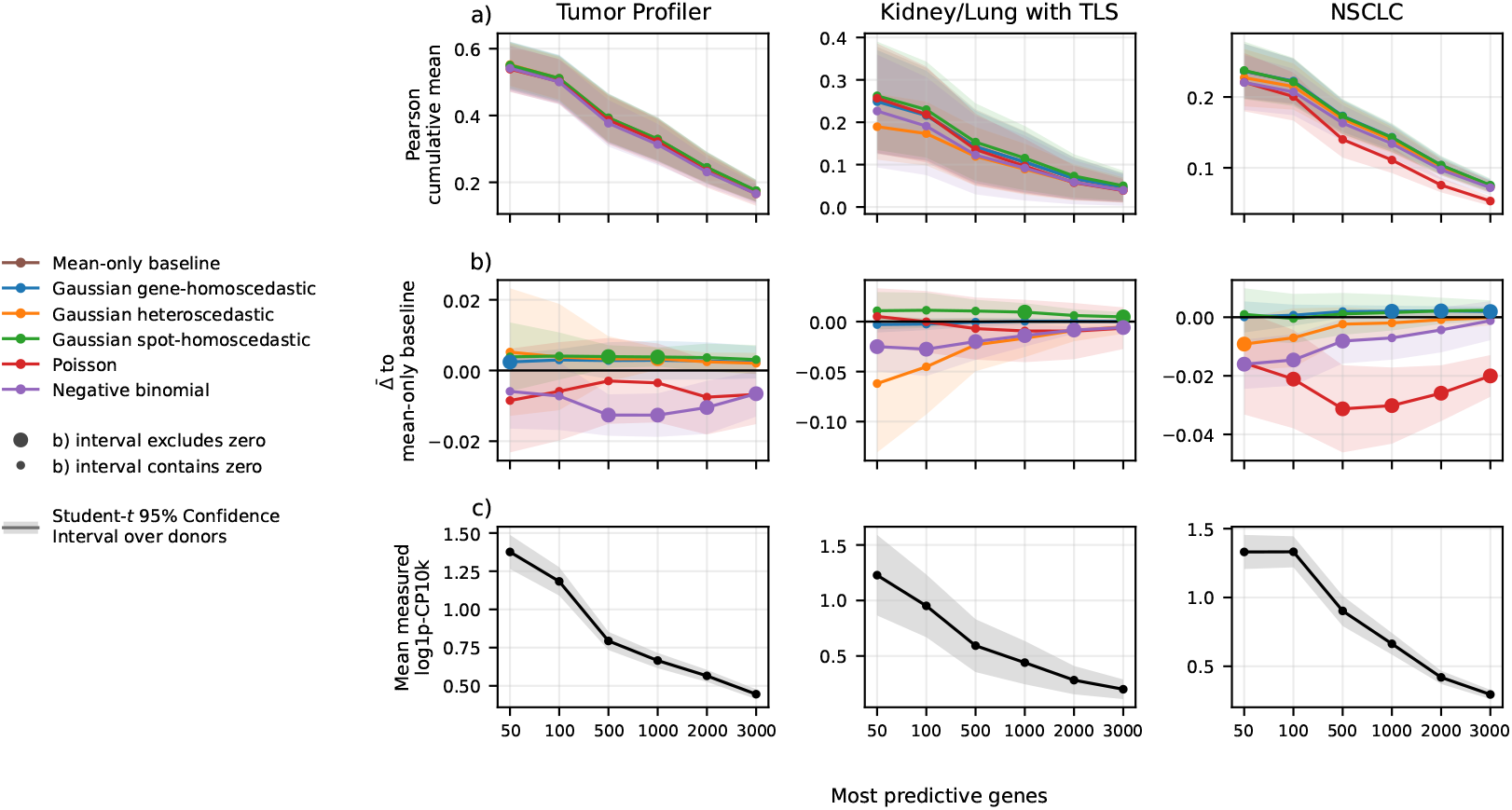
Comparison of the heads’ performance to the mean-only baseline. Genes are ranked once per cohort by their cross-model, donor-averaged correlation. Each point averages the first *n* genes. *a)* Pearson correlation on the log(1 + counts per 10^4^) scale. *b)* Difference to the mean-only baseline taken within each donor. *c)* Model-agnostic mean measured expression.

### A.2 Comparing the two mean estimators

Mapping a count head back to the log(1 + counts per 10^4^) scale admits two estimators (section D): the mean-only *A* and the Jensen-exact *B. B* is used throughout. They differ only by the Jensen gap of the concave log(1 + *·*), so they coincide for a head with no free dispersion and separate as the predictive spread grows. Table 1 shows the difference in correlation values when changing estimators. In the Poisson models, the change of estimator has no impact. In the negative binomial models, estimator *B* results in a higher correlation most of the time. This is particularly the case on the NSCLC cohort.

**Table 1:**
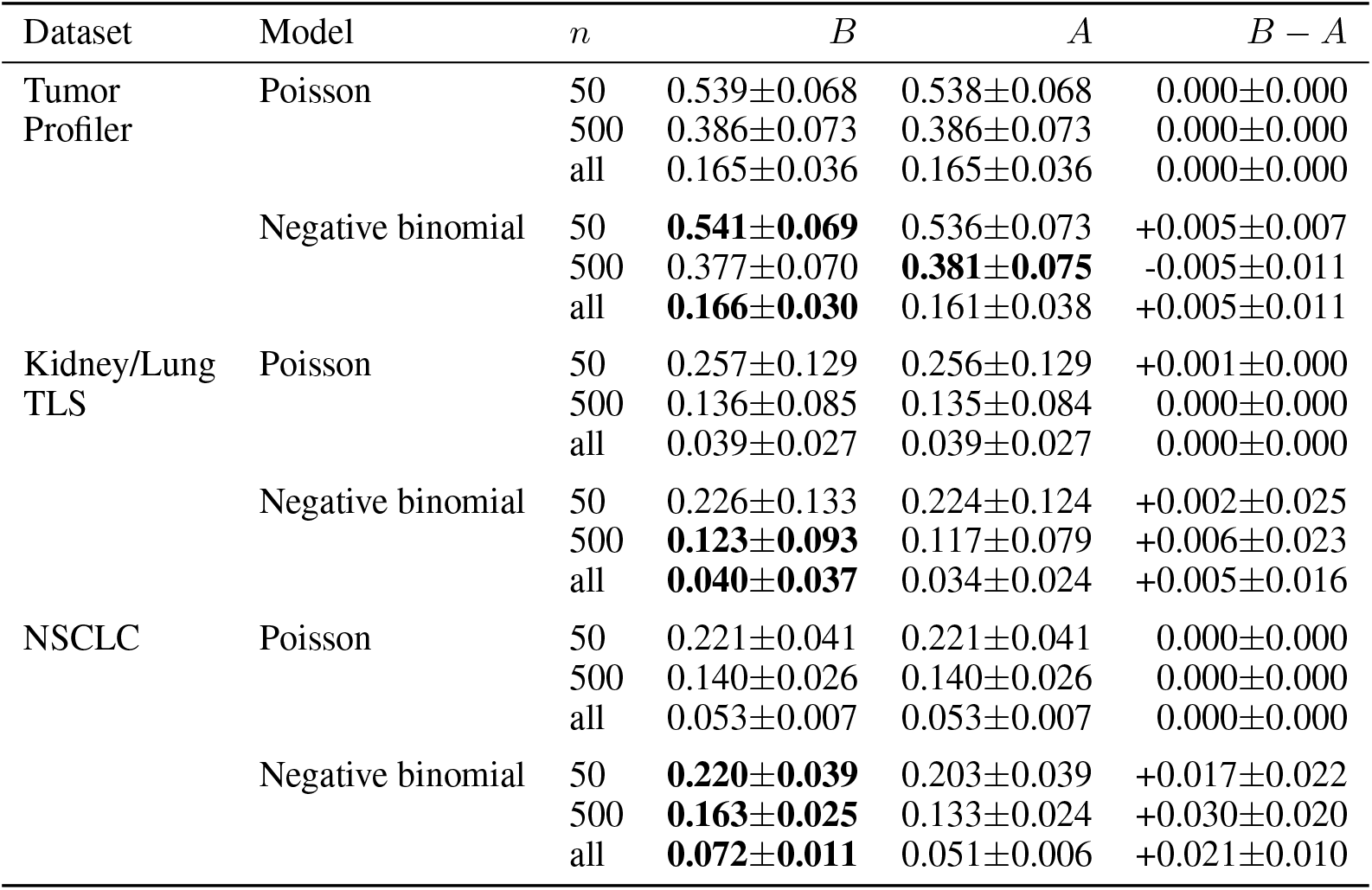
The two estimators of the mean on the log(1 + counts per 10^4^) scale, for the count models. Each entry is the slide-aware Pearson correlation over the first *n* most-predictive genes, averaged over donors. *B* and *A* carry the half-width of the donor Student’s *t* 95% interval on that correlation; *B ™ A* carries it on the per-donor differences. Bold marks the estimator ahead by 0.005 or more at the precision shown.

### A.3 Robustness checks

Section 4.4 reports that the same pattern persists under every modeling and scoring variation we tried. The modeling variations are the epoch selection criterion, the *β*-NLL loss for the Gaussian heads, and training the count heads on raw counts rather than the depth-corrected target. The scoring variations are the absolute log-ratio error in place of the absolute error, and the restriction to the detected entries. Two alternative definitions of the uncertainty are also tested: the coefficient of variation (predicted spread divided by predicted mean) and the dispersion-only quantity 1 + *α*. The latter is the negative binomial predicted variance that would be obtained if every entry were predicted at *µ* = 1. Tables 2 and 3 report the uncertainty-error correlations under those variations, on the gene and spot axes, respectively. A post hoc recalibration of the predicted variance appears there as two further variations; its protocol and its effect on calibration are given in section F.

**Table 2:** Uncertainty-error Spearman correlation across the spots of each gene under each modeling and scoring variation. Correlations are averaged over the held-out donors of the cohort (7, 6, and 8 donors, left to right). Controlled values carry the half-width of the donor Student’s *t* 95% interval. Rows use the reference setting except where stated: absolute error over all entries, the predictive standard deviation as the uncertainty *u*, and the epoch selected on validation correlation. A dash marks a correlation that does not exist.

| Variation | Head | Tumor Profiler |  | Kidney/Lung TLS |  | NSCLC |  |
| --- | --- | --- | --- | --- | --- | --- | --- |
|  |  | uncontrolled | controlled | uncontrolled | controlled | uncontrolled | controlled |
| Reference | Heteroscedastic | +0.20 | +0.06 ± 0.02 | +0.18 | +0.02 ± 0.01 | +0.30 | +0.02 ± 0.01 |
| | Spot-homoscedastic | +0.06 | +0.06 ± 0.01 | −0.15 | −0.01 ± 0.03 | −0.23 | −0.01 ± 0.01 |
|  | Poisson | +0.49 | – | +0.83 | – | +0.76 | – |
| | Negative binomial | +0.36 | +0.03 ± 0.01 | +0.49 | −0.00 ± 0.01 | +0.22 | −0.00 ± 0.01 |
| Detected entries only | Heteroscedastic | +0.11 | +0.12 ± 0.05 | +0.05 | +0.09 ± 0.12 | +0.12 | +0.17 ± 0.04 |
| | Spot-homoscedastic | +0.15 | +0.13 ± 0.04 | +0.10 | +0.04 ± 0.17 | +0.39 | +0.23 ± 0.05 |
|  | Poisson | +0.00 | – | −0.12 | – | −0.20 | – |
| | Negative binomial | +0.06 | +0.09 ± 0.04 | −0.04 | +0.05 ± 0.17 | +0.19 | +0.22 ± 0.05 |
| Absolute log-ratio error | Heteroscedastic | −0.05 | +0.09 ± 0.04 | −0.05 | +0.07 ± 0.10 | −0.02 | +0.15 ± 0.03 |
| | Spot-homoscedastic | +0.08 | +0.09 ± 0.04 | +0.23 | +0.04 ± 0.14 | +0.44 | +0.20 ± 0.04 |
|  | Poisson | −0.26 | – | −0.42 | – | −0.38 | – |
| | Negative binomial | −0.15 | +0.07 ± 0.03 | −0.15 | +0.04 ± 0.17 | +0.09 | +0.21 ± 0.05 |
| Selection: validation NLL | Heteroscedastic | +0.13 | +0.05 ± 0.02 | +0.06 | +0.02 ± 0.01 | +0.11 | +0.00 ± 0.00 |
| | Spot-homoscedastic | +0.08 | +0.05 ± 0.02 | −0.09 | +0.02 ± 0.04 | −0.18 | −0.01 ± 0.01 |
|  | Poisson | +0.49 | – | +0.82 | – | +0.75 | – |
| | Negative binomial | +0.42 | +0.02 ± 0.01 | +0.35 | −0.00 ± 0.01 | +0.34 | +0.00 ± 0.01 |
| Selection: validation MSE | Heteroscedastic | +0.15 | +0.06 ± 0.02 | +0.11 | +0.01 ± 0.01 | +0.21 | +0.01 ± 0.01 |
| | Spot-homoscedastic | +0.06 | +0.05 ± 0.02 | −0.08 | +0.01 ± 0.04 | −0.20 | −0.01 ± 0.01 |
|  | Poisson | +0.49 | – | +0.83 | – | +0.76 | – |
| | Negative binomial | +0.37 | +0.03 ± 0.02 | +0.55 | −0.00 ± 0.00 | +0.34 | −0.00 ± 0.00 |
| β-NLL, $\beta = 0.25$ | Heteroscedastic | +0.19 | +0.06 ± 0.02 | +0.21 | +0.03 ± 0.03 | +0.29 | +0.02 ± 0.01 |
| | Spot-homoscedastic | +0.05 | +0.06 ± 0.02 | −0.15 | +0.00 ± 0.03 | −0.23 | −0.01 ± 0.01 |
| β-NLL, $\beta = 0.5$ | Heteroscedastic | +0.18 | +0.07 ± 0.02 | +0.17 | +0.02 ± 0.03 | +0.29 | +0.02 ± 0.01 |
| | Spot-homoscedastic | +0.05 | +0.06 ± 0.01 | −0.15 | −0.01 ± 0.04 | −0.21 | −0.01 ± 0.01 |
| β-NLL, $\beta = 0.75$ | Heteroscedastic | +0.17 | +0.07 ± 0.02 | +0.16 | +0.02 ± 0.02 | +0.27 | +0.02 ± 0.01 |
| | Spot-homoscedastic | +0.05 | +0.06 ± 0.02 | −0.14 | −0.00 ± 0.04 | −0.23 | −0.00 ± 0.01 |
| β-NLL, $\beta = 1$ | Heteroscedastic | +0.15 | +0.06 ± 0.02 | +0.13 | +0.02 ± 0.03 | +0.22 | +0.02 ± 0.01 |
| | Spot-homoscedastic | +0.04 | +0.06 ± 0.02 | −0.15 | −0.00 ± 0.03 | −0.23 | −0.00 ± 0.01 |
| Raw-count target | Poisson | +0.54 | – | +0.84 | – | +0.82 | – |
| | Negative binomial | +0.54 | +0.02 ± 0.02 | +0.81 | +0.03 ± 0.06 | +0.81 | +0.01 ± 0.01 |
| Recalibrated on $\mu$ | Heteroscedastic | +0.18 | +0.05 ± 0.02 | +0.15 | +0.03 ± 0.02 | +0.14 | +0.01 ± 0.01 |
| | Negative binomial | +0.35 | +0.03 ± 0.01 | +0.41 | −0.00 ± 0.01 | +0.24 | −0.00 ± 0.01 |
| Recalibrated on $u$ | Heteroscedastic | +0.17 | +0.05 ± 0.02 | +0.11 | +0.01 ± 0.01 | +0.16 | +0.01 ± 0.00 |
| | Negative binomial | +0.33 | +0.03 ± 0.01 | +0.41 | −0.00 ± 0.01 | +0.21 | +0.00 ± 0.01 |
| $u = \sqrt{\text{Var}}/\hat{\mu}$ | Gene-homoscedastic | −0.40 | +0.00 ± 0.00 | −0.80 | −0.00 ± 0.00 | −0.73 | −0.00 ± 0.00 |
| | Heteroscedastic | −0.28 | +0.04 ± 0.02 | −0.70 | +0.00 ± 0.01 | −0.65 | +0.01 ± 0.01 |
| | Spot-homoscedastic | −0.35 | +0.04 ± 0.01 | −0.79 | +0.00 ± 0.01 | −0.72 | −0.00 ± 0.01 |
| | Poisson | −0.49 | +0.00 ± 0.00 | −0.83 | +0.00 ± 0.00 | −0.76 | −0.00 ± 0.00 |
| | Negative binomial | −0.23 | +0.03 ± 0.01 | −0.32 | −0.00 ± 0.01 | −0.21 | −0.00 ± 0.00 |
| $u = 1 + \alpha$ | Negative binomial | −0.03 | +0.03 ± 0.01 | −0.24 | −0.00 ± 0.01 | −0.17 | −0.00 ± 0.00 |

**Table 3:** Uncertainty-error Spearman correlation across the genes of each spot under each modeling and scoring variation. Correlations are averaged over the held-out donors of the cohort (7, 6, and 8 donors, left to right). Controlled values carry the half-width of the donor Student’s *t* 95% interval. Rows use the reference setting except where stated: absolute error over all entries, the predictive standard deviation as the uncertainty *u*, and the epoch selected on validation correlation. A dash marks a correlation that does not exist.

| Variation | Head | Tumor Profiler |  | Kidney/Lung TLS |  | NSCLC |  |
| --- | --- | --- | --- | --- | --- | --- | --- |
|  |  | uncontrolled | controlled | uncontrolled | controlled | uncontrolled | controlled |
| Reference | Gene-homoscedastic | +0.34 | -0.02 ± 0.06 | +0.49 | +0.05 ± 0.06 | +0.78 | +0.15 ± 0.04 |
| | Heteroscedastic | +0.66 | +0.15 ± 0.03 | +0.85 | +0.22 ± 0.08 | +0.86 | +0.15 ± 0.02 |
|  | Poisson | +0.79 | - | +0.95 | - | +0.93 | - |
| | Negative binomial | +0.79 | +0.05 ± 0.02 | +0.90 | -0.02 ± 0.04 | +0.92 | -0.00 ± 0.01 |
| Detected entries only | Gene-homoscedastic | -0.09 | -0.07 ± 0.05 | -0.42 | +0.06 ± 0.09 | -0.43 | +0.09 ± 0.04 |
| | Heteroscedastic | -0.02 | -0.01 ± 0.03 | -0.53 | +0.03 ± 0.06 | -0.45 | +0.00 ± 0.03 |
|  | Poisson | +0.34 | - | -0.16 | - | +0.08 | - |
| | Negative binomial | +0.34 | +0.09 ± 0.02 | -0.14 | +0.07 ± 0.04 | +0.09 | +0.06 ± 0.02 |
| Absolute log-ratio error | Gene-homoscedastic | -0.35 | -0.09 ± 0.05 | -0.69 | +0.02 ± 0.11 | -0.78 | +0.03 ± 0.02 |
| | Heteroscedastic | -0.43 | -0.02 ± 0.02 | -0.84 | -0.05 ± 0.08 | -0.79 | -0.10 ± 0.03 |
|  | Poisson | -0.36 | - | -0.85 | - | -0.74 | - |
| | Negative binomial | -0.34 | +0.09 ± 0.02 | -0.81 | +0.07 ± 0.05 | -0.72 | +0.07 ± 0.02 |
| Selection: validation NLL | Gene-homoscedastic | +0.60 | +0.06 ± 0.02 | +0.67 | +0.12 ± 0.06 | +0.81 | +0.18 ± 0.04 |
| | Heteroscedastic | +0.64 | +0.12 ± 0.03 | +0.83 | +0.17 ± 0.09 | +0.86 | +0.11 ± 0.02 |
|  | Poisson | +0.78 | - | +0.94 | - | +0.92 | - |
| | Negative binomial | +0.75 | +0.07 ± 0.03 | +0.93 | -0.01 ± 0.02 | +0.92 | -0.01 ± 0.01 |
| Selection: validation MSE | Gene-homoscedastic | -0.20 | -0.10 ± 0.03 | +0.18 | -0.03 ± 0.04 | +0.49 | +0.02 ± 0.09 |
| | Heteroscedastic | +0.66 | +0.14 ± 0.03 | +0.78 | +0.21 ± 0.10 | +0.86 | +0.15 ± 0.03 |
|  | Poisson | +0.79 | - | +0.95 | - | +0.93 | - |
| | Negative binomial | +0.77 | +0.05 ± 0.04 | +0.89 | -0.01 ± 0.04 | +0.92 | +0.01 ± 0.02 |
| β-NLL, $\beta = 0.25$ | Gene-homoscedastic | +0.16 | -0.04 ± 0.04 | +0.49 | +0.07 ± 0.07 | +0.76 | +0.14 ± 0.06 |
| | Heteroscedastic | +0.65 | +0.14 ± 0.03 | +0.83 | +0.25 ± 0.10 | +0.86 | +0.15 ± 0.02 |
| β-NLL, $\beta = 0.5$ | Gene-homoscedastic | +0.26 | -0.02 ± 0.04 | +0.48 | +0.05 ± 0.05 | +0.78 | +0.15 ± 0.04 |
| | Heteroscedastic | +0.66 | +0.15 ± 0.03 | +0.82 | +0.26 ± 0.08 | +0.86 | +0.17 ± 0.02 |
| β-NLL, $\beta = 0.75$ | Gene-homoscedastic | +0.31 | -0.01 ± 0.04 | +0.47 | +0.05 ± 0.04 | +0.77 | +0.14 ± 0.05 |
| | Heteroscedastic | +0.65 | +0.14 ± 0.04 | +0.80 | +0.25 ± 0.06 | +0.85 | +0.19 ± 0.02 |
| β-NLL, $\beta = 1$ | Gene-homoscedastic | +0.27 | -0.02 ± 0.04 | +0.56 | +0.07 ± 0.02 | +0.75 | +0.13 ± 0.05 |
| | Heteroscedastic | +0.65 | +0.14 ± 0.04 | +0.77 | +0.24 ± 0.04 | +0.83 | +0.21 ± 0.03 |
| Raw-count target | Poisson | +0.81 | - | +0.94 | - | +0.93 | - |
| | Negative binomial | +0.81 | +0.07 ± 0.03 | +0.90 | -0.02 ± 0.02 | +0.93 | +0.01 ± 0.02 |
| Recalibrated on $\mu$ | Heteroscedastic | +0.62 | +0.15 ± 0.06 | +0.72 | +0.14 ± 0.10 | +0.80 | +0.12 ± 0.03 |
| | Negative binomial | +0.74 | +0.04 ± 0.02 | +0.67 | -0.05 ± 0.05 | +0.86 | -0.02 ± 0.01 |
| Recalibrated on $u$ | Heteroscedastic | +0.62 | +0.15 ± 0.05 | +0.73 | +0.16 ± 0.11 | +0.81 | +0.13 ± 0.03 |
| | Negative binomial | +0.73 | +0.04 ± 0.02 | +0.60 | -0.05 ± 0.06 | +0.83 | -0.01 ± 0.02 |
| $u = \sqrt{\text{Var}}/\hat{\mu}$ | Gene-homoscedastic | -0.60 | -0.03 ± 0.01 | -0.91 | +0.02 ± 0.02 | -0.88 | +0.05 ± 0.02 |
| | Heteroscedastic | -0.60 | +0.07 ± 0.03 | -0.83 | +0.08 ± 0.03 | -0.84 | +0.08 ± 0.01 |
| | Spot-homoscedastic | -0.61 | +0.00 ± 0.00 | -0.90 | +0.00 ± 0.00 | -0.88 | -0.00 ± 0.00 |
| | Poisson | -0.79 | -0.00 ± 0.00 | -0.95 | +0.00 ± 0.00 | -0.93 | -0.00 ± 0.00 |
| | Negative binomial | -0.77 | +0.07 ± 0.02 | -0.89 | +0.03 ± 0.04 | -0.92 | +0.01 ± 0.01 |
| $u = 1 + \alpha$ | Negative binomial | -0.69 | +0.06 ± 0.02 | -0.62 | +0.05 ± 0.08 | -0.91 | +0.01 ± 0.01 |

### A.4 Uncertainty-error correlation for the other sources

The body reports the head’s own predicted spread. The same measurement was repeated on the epistemic term of the three estimators and on the training-free neighborhood signal, and on the STimage readout, scored identically on all entries (figs. 5 to 7).

**Figure 5:**
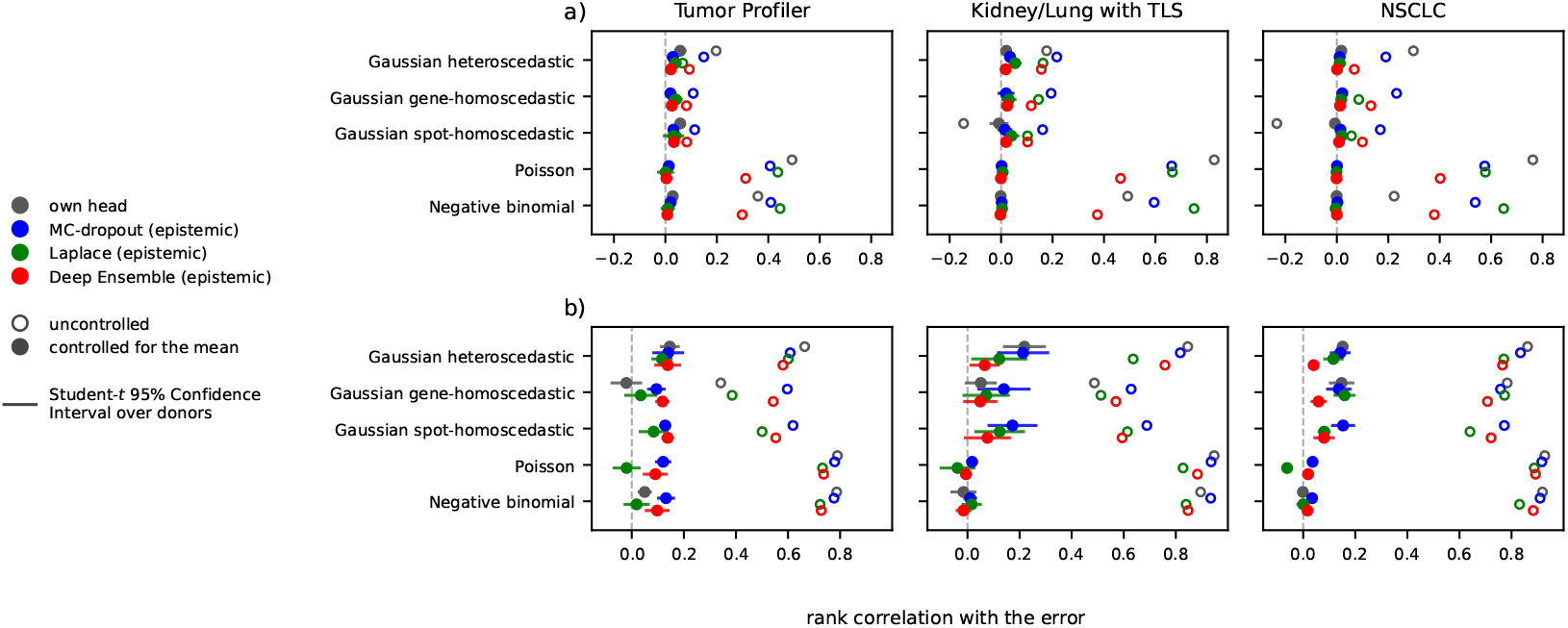
Uncertainty-error Spearman correlation for four sources of uncertainty, uncontrolled and controlling for the predicted mean, across the spots of one gene (top row) and across the genes of one spot (bottom row).

**Figure 6:**
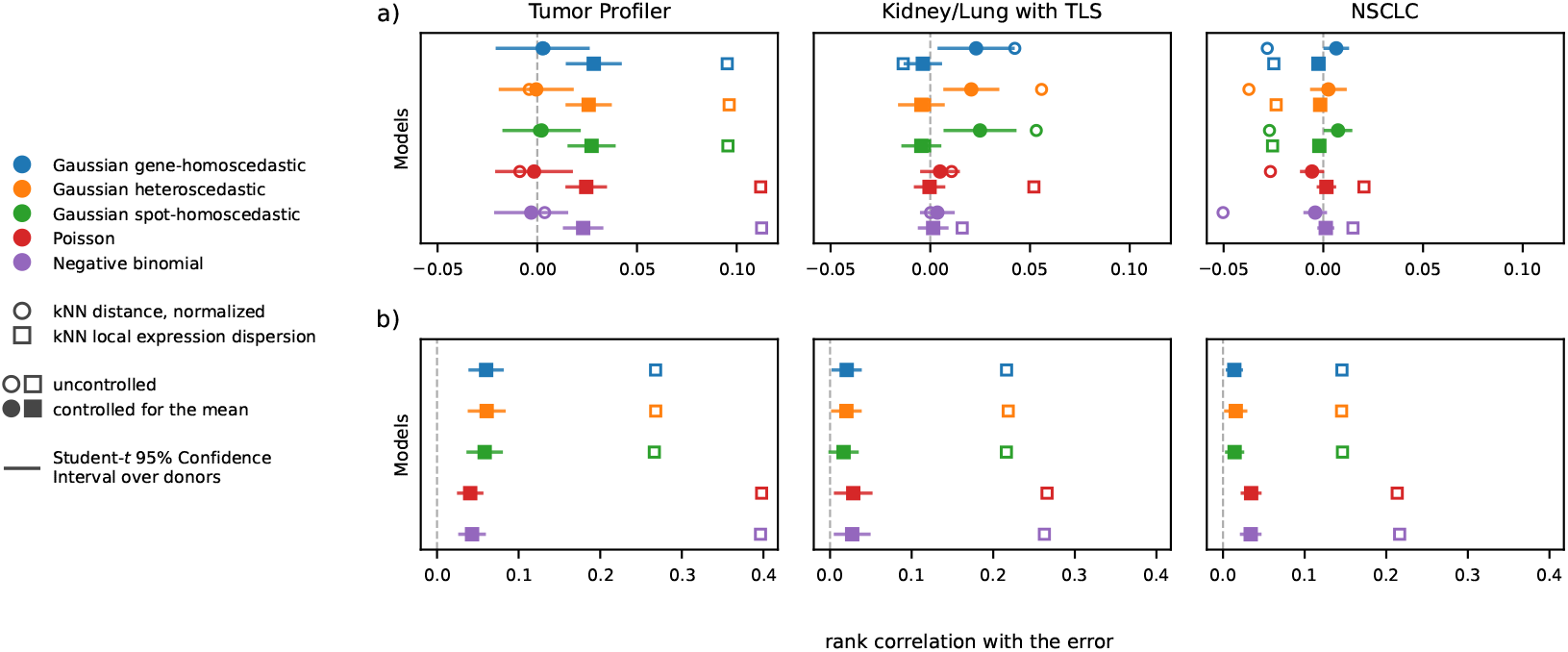
Uncertainty-error Spearman correlation for two training-free neighborhood signals, uncontrolled and controlling for the predicted mean, across the spots of one gene (top row) and across the genes of one spot (bottom row). Circle, the mean distance to a spot’s nearest neighbors divided by the radius of the cluster it falls in; square, the dispersion of the expression measured among those neighbors. Colors are the predictive heads, which supply only the error being ranked. The distance takes one value per spot and is therefore absent from the bottom row.

**Figure 7:**
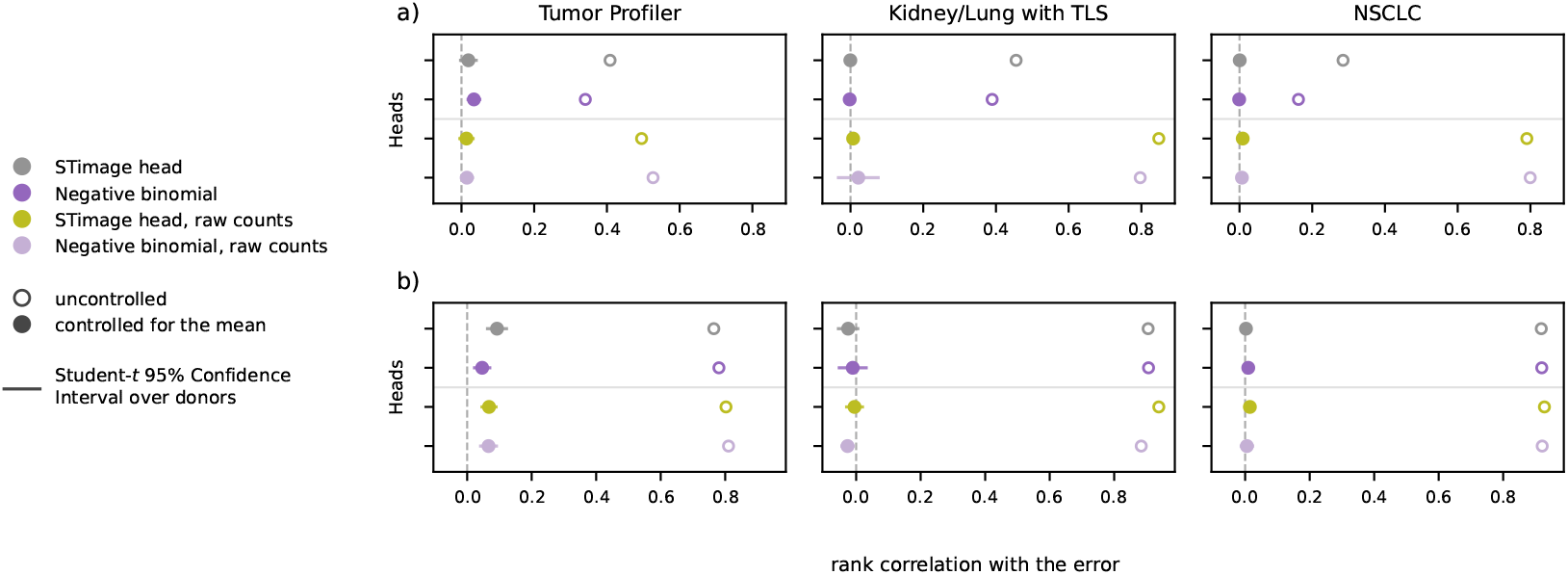
Uncertainty-error Spearman correlation for the STimage readout and our negative binomial head, uncontrolled and controlling for the predicted mean, across the spots of one gene (top row) and across the genes of one spot (bottom row).

No epistemic term separates itself from the aleatoric one: across the spots of one gene every value stays below +0.06, and which of the two is higher changes with the cohort. The epistemic term is defined even where the head’s own spread carries no signal, for the gene-homoscedastic Gaussian and for the Poisson.

Figure 6 reports the training-free neighborhood signal, computed from the frozen embeddings and the measured expression alone, so a head enters it only through the error being ranked. Across the genes of one spot the dispersion term reaches +0.01 to +0.06, below the +0.15 to +0.22 the heteroscedastic Gaussian reaches on the same grouping. Across the spots of one gene the fitted variance is the higher of the two in all three cohorts. The distance term stays below +0.025, and it is absent across the genes of one spot, taking a single value per spot.

The STimage readout was retrained under our embeddings, splits, and scoring, on both count targets, and scored exactly as our own heads (fig. 7). Its uncertainty-error correlations are indistinguishable from ours.

### A.5 Concentration on the worst-predicted genes

The numbers in section 4.3 are averages over genes, so we ask whether individual genes depart from that average (fig. 8). The highest controlled correlation between uncertainty and error is found on the worst-predicted genes. For each of the three cohorts and each of the three models measurable on this grouping, the 100 genes carrying the strongest controlled correlation systematically have a lower median accuracy than the panel as a whole. On NSCLC their median accuracy is 0.02 against 0.07 over all genes. The uncertainty is least informative on the well-predicted genes, which are the ones a user would rely on. The same concentration appears for the epistemic term and for the training-free neighborhood signal.

**Figure 8:**
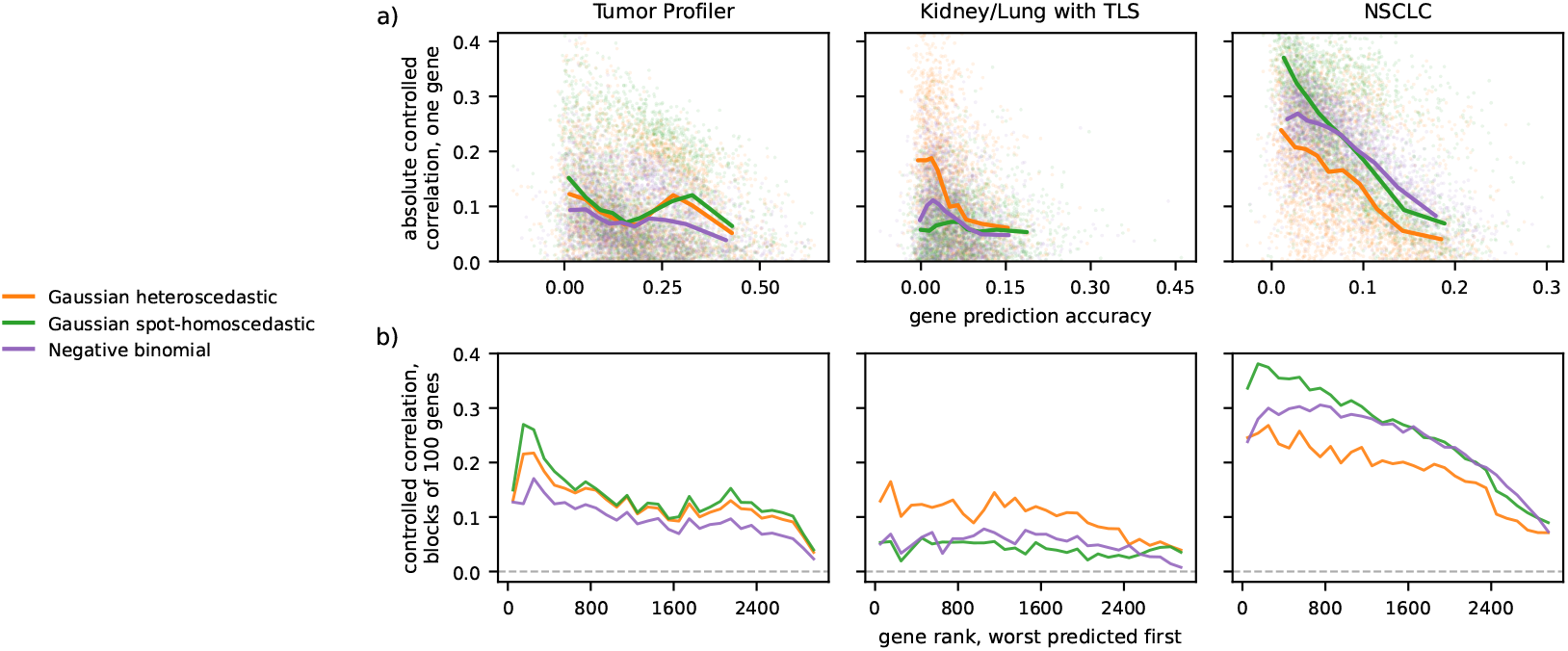
Controlled uncertainty-error correlation on the gene axis, against gene predictability. *a)* one point per gene, accuracy against the absolute controlled correlation, with the median over accuracy deciles curve drawn on top. *b)* same correlation along gene accuracy ordering.

### A.6 Sparsification curves

Figures 9 and 10 report the curves on all entries and on the detected entries, respectively, and table 4 the share of the oracle each score recovers. On all entries, withholding within a (gene, slide) group, the predicted mean used alone is as good a withholding score as the predicted uncertainty, or better: at an undetected entry, the error is the predicted mean itself, so ranking by the mean partly ranks by the error. Restricting to the detected entries removes that mechanism. Withholding at a fixed predicted mean, the non-Poisson heads then beat the mean alone in almost every cohort, but recover at most 30% of what the oracle achieves, which lowers the mean error by at most 16% when half of the predictions are withheld. Without the mean bins, the mean alone remains a strong score: the spot-homoscedastic Gaussian reaches the largest share of any head, 44% on NSCLC, but the predicted mean alone already reaches 33% there, and this relatively high value is not reproduced on the other two cohorts.

**Table 4:** Share of the oracle’s error reduction recovered by the predicted uncertainty (*u*) and by the predicted mean used alone (*µ*), as defined in section E. A share of 0 is an uninformative ranking, 1 matches the oracle. The *µ* column takes the better of the two withholding directions: the largest predicted means first on all entries, the smallest predicted means first on the detected entries, except where marked *†*. The Poisson uncertainty is a monotone function of its mean, so *u* is exactly the largest-means-first ranking and the two columns coincide wherever that is also the better *µ* direction.

| Setting | Head | Tumor Profiler |  | Kidney/Lung TLS |  | NSCLC |  |
| --- | --- | --- | --- | --- | --- | --- | --- |
| | | $u$ | $\mu$ | $u$ | $\mu$ | $u$ | $\mu$ |
| All entries<br>within (gene, slide) | Heteroscedastic | +0.21 | +0.20 | +0.13 | +0.28 | +0.18 | +0.28 |
|  | Spot-homoscedastic | +0.12 | +0.19 | −0.07 | +0.26 | −0.13 | +0.29 |
|  | Poisson | +0.34 | +0.34 | +0.28 | +0.28 | +0.27 | +0.27 |
|  | Negative binomial | +0.33 | +0.35 | +0.17 | +0.24 | +0.15 | +0.27 |
| All entries<br>at fixed mean | Heteroscedastic | +0.09 | +0.04 | +0.04 | +0.06 | +0.06 | +0.06 |
|  | Spot-homoscedastic | +0.08 | +0.04 | +0.02 | +0.06 | −0.02 | +0.06 |
|  | Poisson | +0.08 | +0.08 | +0.06 | +0.06 | +0.06 | +0.06 |
|  | Negative binomial | +0.11 | +0.09 | +0.04 | +0.05 | +0.05 | +0.06 |
| Detected entries<br>within (gene, slide) | Heteroscedastic | +0.13 | −0.02 | +0.12 | +0.18 | +0.13 | +0.31 |
|  | Spot-homoscedastic | +0.17 | −0.01 | −0.01 | +0.20 | +0.44 | +0.33 |
|  | Poisson | +0.11 | +0.11† | −0.02 | +0.01 | −0.06 | +0.03 |
|  | Negative binomial | +0.20 | +0.11† | +0.12 | +0.05 | +0.33 | +0.06 |
| Detected entries<br>at fixed mean | Heteroscedastic | +0.11 | +0.01 | +0.17 | +0.04 | +0.16 | +0.08 |
|  | Spot-homoscedastic | +0.12 | +0.01 | −0.00 | +0.05 | +0.28 | +0.09 |
|  | Poisson | +0.03 | +0.03† | +0.00 | +0.00 | −0.01 | +0.01 |
|  | Negative binomial | +0.13 | +0.03† | +0.16 | +0.01 | +0.30 | +0.01 |

**Figure 9:**
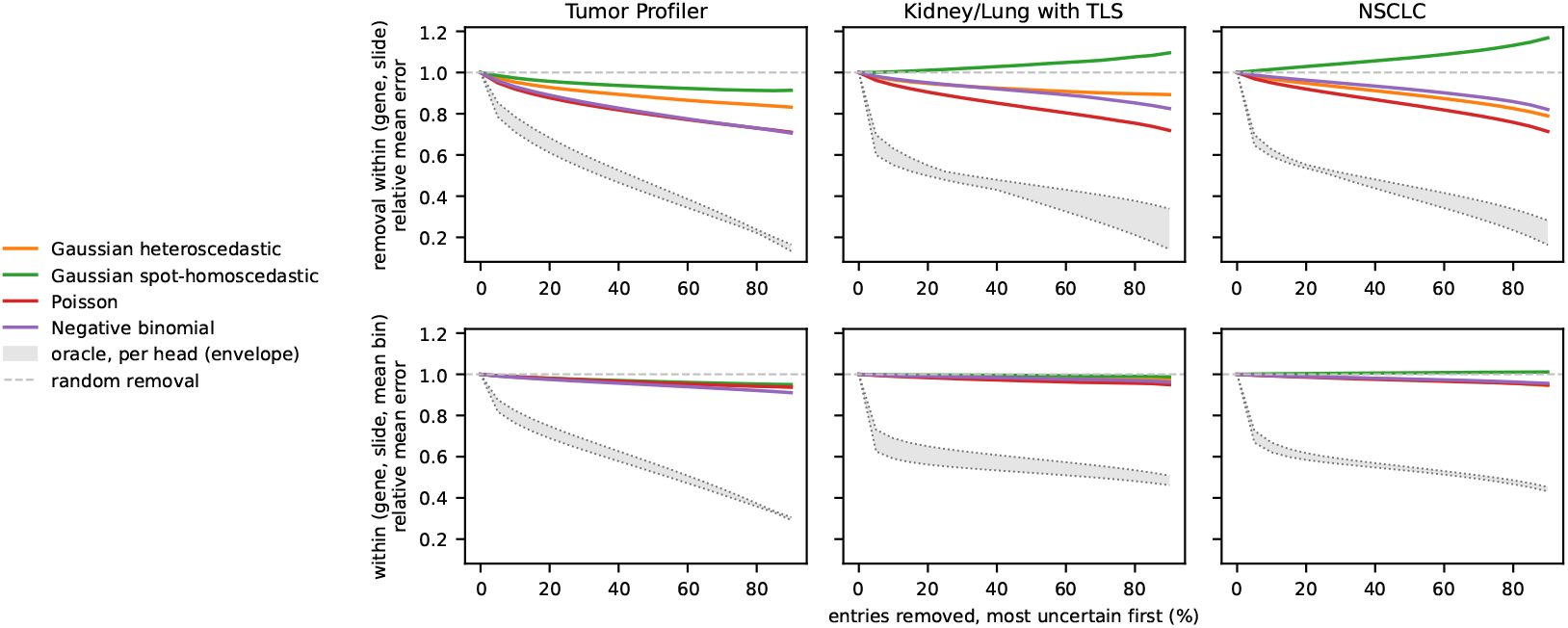
Sparsification curves on all entries, withholding within a gene and slide (top) and within a bin of the predicted mean of that group (bottom). Gray band, the oracle of each head; dashed line, withholding at random.

**Figure 10:**
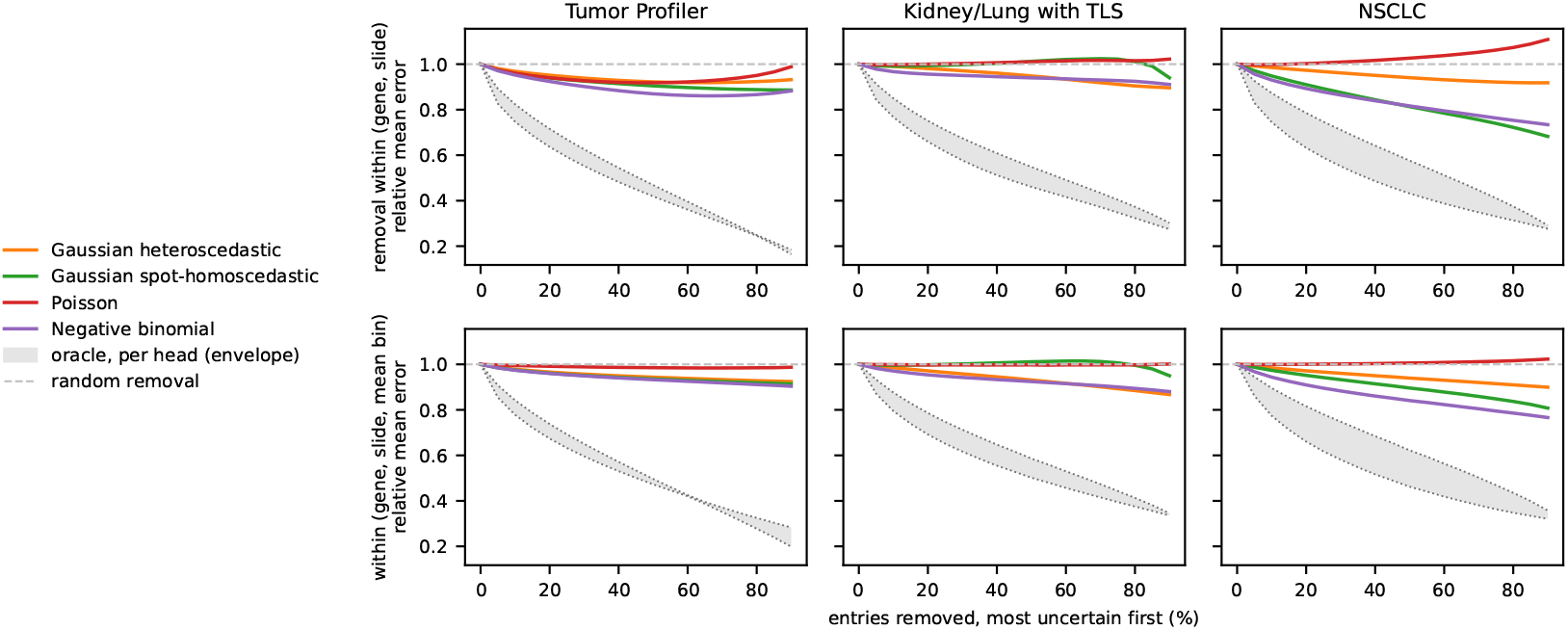
Sparsification curves on the detected entries, with the same two protocols and reference levels as fig. 9.

### A.7 Group-level structure and behavior under distribution shift

We show that a spread that fails to order the errors inside a group can still be structured. At a matched predicted mean, obtained by pairing spots inside one gene and one slide on equal predicted means, the non-tumor tissue types carry more uncertainty than the tumor (fig. 11): *×*1.23 on Tumor Profiler and *×*1.16 on Kidney/Lung TLS. The effect is not gene-specific: the head predicting one variance per spot, shared by all 3,000 genes, returns *×*1.22.

**Figure 11:**
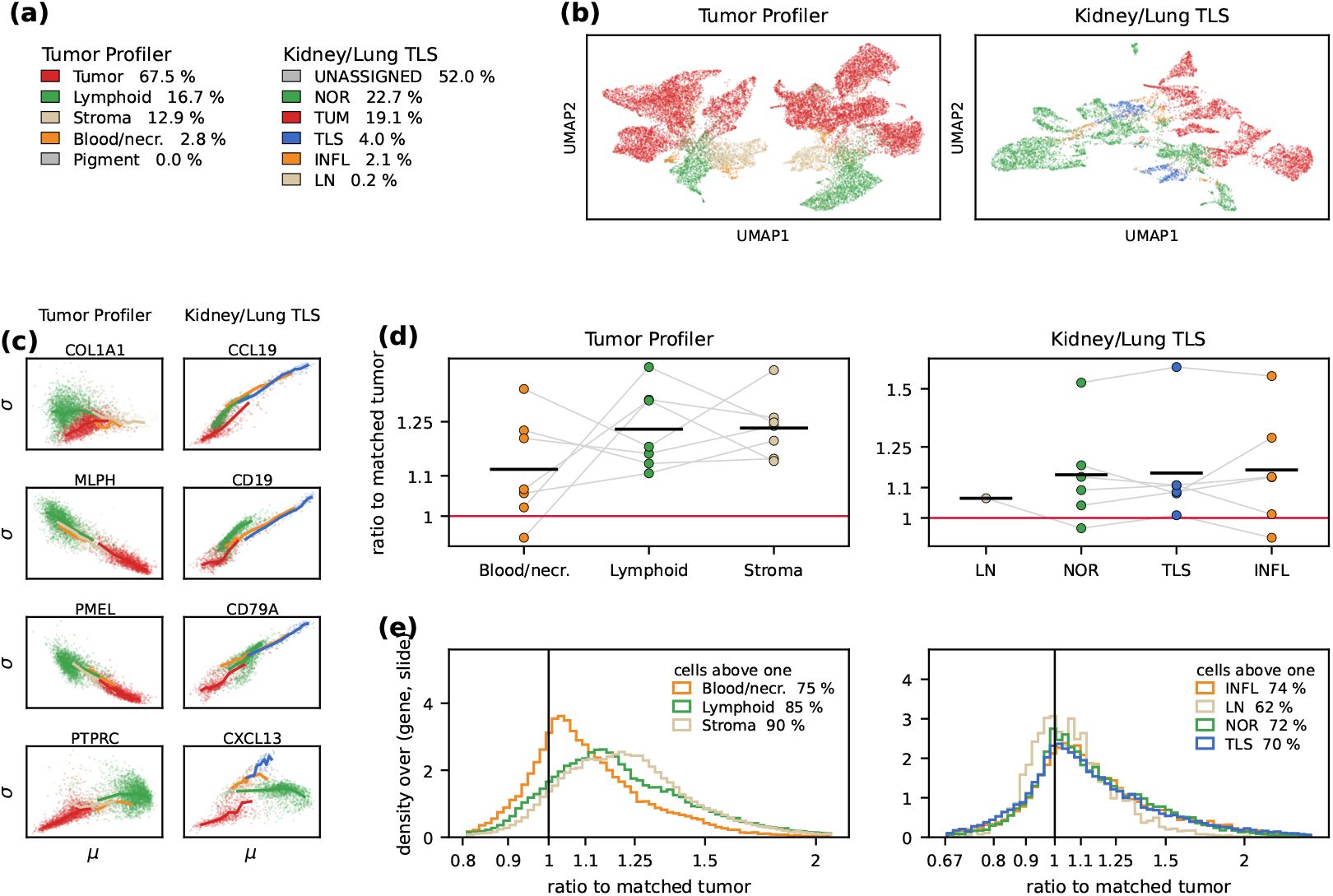
Predicted uncertainty by tissue type after matching on predicted mean. Tumor Profiler on the left of every block, Kidney/Lung TLS on the right. *a)* tissue types and their share of spots, gray for the categories excluded. *b)* UMAP of the frozen patch representations, colored by tissue type. *c)* predicted uncertainty against predicted mean on one slide, one point per spot, one gene per square; lines are medians inside ten equal-count bins of the predicted mean. *d)* ratio to the matched tumor spot, one point per donor, gray lines joining the tissue types of one donor, red line at equality. *e)* the same ratio over every (gene, slide) pair.

#### Matching on the predicted mean

Spots are paired inside a (gene, slide) group, so neither gene identity nor slide can contribute to the contrast. Every annotated spot is matched to the tumor spot of that group whose predicted mean is closest, greedily and with replacement, and a pair is kept only when the two predicted means differ by less than one tenth of the standard deviation of the predicted mean over the annotated spots of that group. Spots left without a partner are dropped; the share finding one has a median between 0.95 and 1.00 across cells. A (gene, slide, tissue type) cell is retained at ten pairs or more. Cells are averaged inside each held-out donor, and the *K* donor values give the reported ratio and its interval. On Kidney/Lung TLS the unassigned spots, about half of the cohort, are excluded, and the lymph-node category is 33 spots on a single slide.

The two uncertainty components respond differently when the input leaves the training distribution. Under a Gaussian blur applied before patch extraction, the slide-aware correlation collapses, from 0.393 to 0.078 on Tumor Profiler, and the embedding moves far from the training pool. The aleatoric term *decreases* as the model gets worse, Monte Carlo dropout stays flat, and only the distance-based and deep ensemble epistemic terms grow [46]. Detecting inputs that have left the training distribution is the clearest use case supported by these estimates, and it is a different question from ranking errors at a fixed distribution, which is the question addressed in the body. The same pattern is shown on NSCLC in fig. 12, and the other cohorts behave the same way.

**Figure 12:**
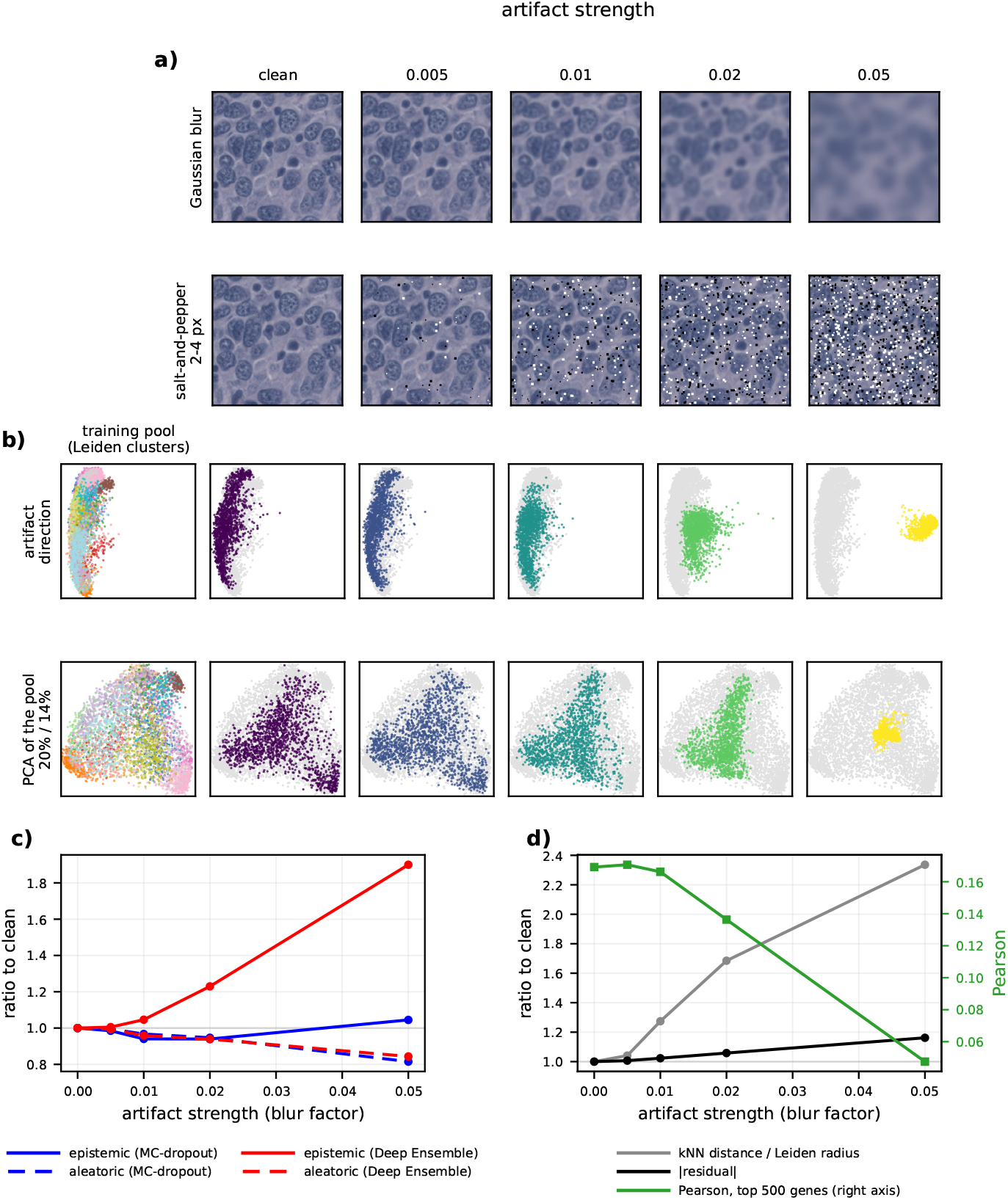
Effect of a synthetic image artifact on prediction accuracy and uncertainties. On NSCLC, artifact strength increases to the right. *a)* one Visium spot at each level of the two artifacts, Gaussian blur and salt-and-pepper noise. *b)* the same patient at each blur level, represented on a plane whose first axis is the mean displacement the artifact causes, and, below, on the ordinary PCA plane of the training pool; first column the pool alone, colored by its Leiden clusters, elsewhere the pool is gray. Patient color indicates artifact severity. *c)* predicted uncertainty relative to its clean value. *d)* distance to the training pool and absolute error on the same scale, with the slide-aware Pearson correlation on the right axis. In *c)* and *d)* every quantity carrying a gene axis is restricted to the 500 best-predicted genes and averaged over held-out donors.

#### Synthetic artifacts

Two artifacts are applied to the slides before patch extraction: a Gaussian blur whose standard deviation is proportional to the spot diameter and salt-and-pepper noise on a fraction of the pixels of the model input, each at increasing severity. The models are frozen and prediction is repeated on the modified embeddings. The artifact does reach the model: at the strongest blur, a spot’s embedding has a cosine similarity of 0.03 to 0.12 with its clean one, and its distance to the training pool is multiplied by 1.7 to 2.3.

## B Data and preprocessing

### Cohorts

Tumor Profiler: 18 FFPE sections from 7 patients with metastatic melanoma, with expert spot annotations (Tumor, Normal lymphoid tissue, Blood/Necrosis, Stroma, Pigment) [43]. Kidney/Lung TLS: FFPE sections from kidney and lung tumors enriched for tertiary lymphoid structures, with expert annotations (TLS, Immune, Tumor, Normal, Lymph node, Unassigned) [44]; one lung and one kidney slide were excluded for lower quality following Nonchev et al. [4], leaving 6 sections from 6 patients. NSCLC: 36 fresh-frozen sections from 8 patients, 20 tumor and 16 matched adjacent non-involved, pooling 5 lung adenocarcinoma and 3 lung squamous cell carcinoma patients into a single cohort [45]. All three use the 10x Visium assay, with 77,338, 19,437, and 68,659 spots, respectively. Kidney/Lung TLS is publicly available on Zenodo [44] and NSCLC from BioStudies/ArrayExpress under accession E-MTAB-13530; Tumor Profiler is accessible upon request through the Tumor Profiler Consortium’s portal at https://tumorprofilercenter.ch/contacts.

### Patch extraction and embedding

Each spot’s input is a square patch centered on the spot, with a side of 1.5 spot diameters (82.5 *µ*m for the 55 *µ*m capture spot), following ST-Net [2]; restricting the patch to the spot diameter was verified to perform worse. The half-width is computed in each slide’s own pixel scale, so the physical field of view is constant across slides scanned at different magnifications. Patches are resized to 224 × 224 and embedded with Midnight-12k [32], frozen. The spot representation is the class token concatenated with the mean of the 256 patch tokens, giving the 3,072-dimensional vector *x*_*s*_.

### Expression targets

Writing *y*_*sg*_ for the raw count and *ℓ*_*s*_ = 10^−4^ Σ_*g*_ *y*_*sg*_ for the library size, the Gaussian heads are trained on log(1 + *y*_*sg*_*/ℓ*_*s*_). The count heads place mass only on integers, so they are trained on round 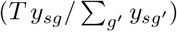, every spot placed at a common depth *T* rather than its own, so that no per-spot library size enters the prediction. We set *T* = 25,000 for Tumor Profiler and *T* = 10,000 for the two others. The choice trades two errors: too low a *T* rounds small non-zero counts to zero, too high a *T* asserts a sampling precision the data do not have. At these depths, rounding sends 1.6%, 8.7%, and 2.5% of the non-zero entries to zero, against 57.5% and 28.1% on Kidney/Lung TLS and NSCLC had *T* been set at each cohort’s median library size. Each cohort is restricted to its 3,000 most highly variable genes, selected with the seurat_v3 procedure [19] applied to all spots at once with the slide as batch key.

### The cost of rounding

Every count head was also trained on the same target without rounding, so that only the rounding differs between the two runs. The two versions reach the same performance at the depths used here, with no per-donor interval excluding zero. A cost appears only when *T* is lowered to each cohort’s median library size on Kidney/Lung TLS and NSCLC, and it grows with the share of non-zero values the rounding destroys (fig. 13).

**Figure 13:**
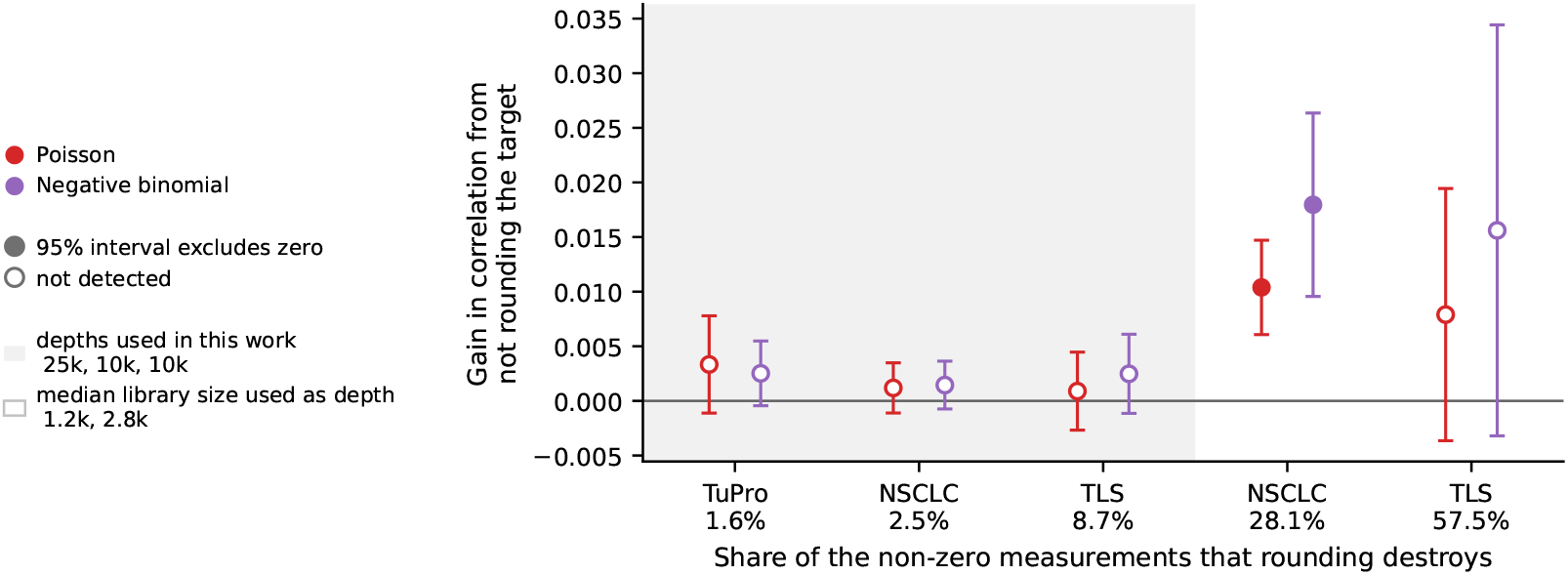
Gain in correlation from training the count heads on the unrounded target, against the share of non-zero measurements that rounding destroys at that depth. The shaded band marks the depths used in this work; the unshaded conditions place each cohort at its median library size. The depth used for Tumor Profiler is its median library size.

## C Model, training, and architecture search

### Trunk and heads

The trunk is a two-layer perceptron, each layer a linear map followed by layer normalization, ReLU, and dropout 0.1, with widths (1024, 256). Every parameter of the prediction is an affine readout of that representation with one channel per gene; the parameters constrained to be positive are read out in log space and exponentiated. The objective is the negative log-likelihood averaged over spots and over genes, so every gene carries the same weight and the magnitude does not depend on the panel size.

### Optimization and folds

Adam, batch size 512, learning rate 8 *×* 10^−4^ ramped linearly over three epochs, at most 25 epochs. The retained epoch maximizes the slide-aware Pearson correlation on the validation donor, evaluated through the mean only. No model is selected on its calibration or on its likelihood; changing the selection criterion to MSE or the negative log-likelihood was found to reduce mean performance, while the uncertainty still failed to track errors. Under leave-one-patient-out, each donor is the test set exactly once, the next donor in a fixed ordering is the validation set, and the remaining *K ™* 2 donors are the training set. Since the folds share most of their training donors, the Student’s *t* intervals over donors are optimistic approximations.

### Architecture search

The trunk was tuned once, on the mean-only baseline as a head-agnostic proxy, and then used unchanged for every cohort and likelihood. Seven axes were searched: hidden dimensions, dropout, activation, weight decay, base learning rate, batch size, and a normalization toggle. From a grid of 2,304 combinations, 256 were drawn at random under a fixed seed. Learning rate and batch size were coupled by square-root scaling, so that a large batch is judged on whether it is viable rather than penalized for being under-tuned. BatchNorm was excluded by design: it would make a spot’s prediction, and hence its predicted uncertainty, depend on the other spots in its mini-batch. The search ran on Tumor Profiler, the highest-quality cohort, over five grouped donor folds.

### Architecture is not the binding constraint

Among the non-degenerate configurations the score spans *≈*0.024 in Pearson correlation, no larger than the fold-to-fold standard deviation of a single configuration (*≈*0.019), so the leading configurations are indistinguishable. The single failure mode is strong weight decay combined with no normalization or a high learning rate, and that degenerate tail alone accounts for the full best-to-worst band of 0.140. Layer normalization did not measurably change accuracy (*≈*0.003) but removed that failure mode and roughly halved the across-configuration variance. Re-running the search on the two other cohorts left the per-axis structure unchanged, the Tumor Profiler configuration ranking 21st and 10th of 256, respectively. This is why the negative result is not attributed to model capacity. We attribute differences in mean performance between cohorts to cohort quality and signal strength rather than architecture.

### The STimage readout

STimage fits a negative binomial per spot and gene on top of a frozen convolutional backbone, its only trainable weights being a dense layer of two units per gene [14].^6^ That head is reproduced here and fed the same Midnight embedding, so it reads out of *x*_*s*_ directly rather than of the trunk, for a gene *g* and spot *s*,

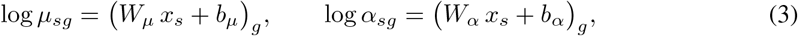

resulting in 18.4 million parameters versus 5.0 million for the negative binomial head. Its published recipe is reproduced as well: learning rate 10^−4^, batches of 128 spots, at most 100 epochs with patience 10, and the epoch selected on the validation negative log-likelihood. The likelihood, the target, the embeddings, the folds, and the scoring are those of every other head; the optimizer and the selection criterion are STimage’s own.

### Compute

All models were trained on a shared cluster, one NVIDIA GPU per job, predominantly a GeForce RTX 4090. Every reported number is a leave-one-patient-out cross-validation, that is six to eight models per configuration and cohort. The configurations run are: mean-only, Gaussian spot-homoscedastic, gene-homoscedastic, and heteroscedastic, negative binomial and Poisson on both the depth-corrected and the raw count targets, and the STimage readout. Each Gaussian is trained once per beta value, a sweep the count heads do not have. The ensembles train *M* = 5 members per configuration, for eight of these heads and for the STimage readout. The trunk being a small multilayer perceptron over frozen embeddings, a single cross-validation takes about five minutes on the GPU, each fold stopping on a validation criterion with a patience of five epochs and running 16 epochs on average against a ceiling of 25. The STimage readout was trained under its own published recipe, which gives it a longer schedule, about twenty minutes per cross-validation at 24 epochs on average. Monte Carlo dropout, the Laplace approximation, and the neighborhood signal are computed post hoc and train nothing.

## D Uncertainty estimators

### Two estimators on the common axis

A count head predicts integer counts at depth *T* , so mapping it to log(1 + counts per 10^4^) is not unique. With *δ* = *T/*10^4^, the mean-only estimator is *A* = log(1 + *µ/δ*) and the predictive mean of the transform is *B* = E[log(1 + *Y/δ*)] = _*k≥*0_ log(1 + *k/δ*) *P* (*Y* = *k*). They differ by the Jensen gap of the concave log(1 + *·*), which is not a constant but grows with the predictive spread, so a head with a free dispersion is exactly the one for which the choice matters. *A* answers whether the mean prediction is degraded; *B* is the exact analogue of the estimand of a Gaussian trained on that scale. Estimator *B* was found to give a better mean performance on the validation donor, and is used throughout. The two estimators are compared in section A.2.

### Epistemic estimates

Deep ensembles use *M* = 5 members differing only in their random seed, each a complete run trained and early-stopped on its own [25]. Monte Carlo dropout reuses each trained checkpoint with 30 forward passes at the training dropout rate [26]. The Laplace approximation is applied to the last layer only [27], with a per-gene Gauss-Newton posterior whose prior precision is selected per gene on a grid by the predictive negative log-likelihood on the validation donor; for the count heads the exponential link makes the mean lognormal under the posterior, with moments in closed form, so the predictive is never sampled. For the Laplace count heads, the posterior variance is large enough on a small fraction of entries for that lognormal mean to exceed any plausible target value, reaching 10^3^ or more on fewer than 0.10% of entries against a median of 0.2. Rank correlations are unaffected, but mean-based statistics are, so these Laplace count heads are excluded from the sparsification analysis. All three epistemic estimators combine with the aleatoric part through the law of total variance,

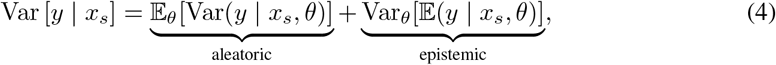

with *θ* the parameters under the posterior each estimator defines, evaluated on the scale the model was trained on.

### Neighborhood signal

Embeddings are L2-normalized and the reference pool for a patient is formed by the other patients of the same cohort [13]. Neighbors are weighted by a Gaussian kernel of their distance and results are reported at *k* = 25. The aleatoric analogue is the weighted standard deviation of the expression among the *detected* neighbors only, since a spread taken over zeros and detected values together would mainly report the detection rate; it is left undefined below five detected neighbors. The epistemic analogue is the mean distance to the neighbors, divided by the radius of the cluster the spot falls in, so that a rare spot type does not read as an unfamiliar morphology. Clusters are defined by Leiden clustering with a resolution of 1.

## E Metrics

### Accuracy

A slide on which prediction or target is constant leaves the slide-aware Pearson correlation undefined and is excluded rather than counted as zero. Curves along the gene axis use one ordering per cohort, from the best to the worst predicted, fixed once and shared by every curve of that cohort, so a given *n* denotes the same gene set everywhere.

### Calibration

For an observation *y* with predictive CDF *F* , the probability integral transform *F* (*y*) is uniform on [0, 1] under a correct continuous predictive. A discrete predictive cannot reach that uniformity, its CDF being a step function. Thus, we use the randomized transform *F* (*y*^−^) + *U* [*F* (*y*) *™ F* (*y*^−^)], with *U ∼* Uniform(0, 1), and replace *U* by its expectation, giving the seed-free nonrandomized form of Czado et al. [33]. Writing *F*_lo_(*y*) = ℙ(*Y < y*), *F*_hi_(*y*) = ℙ(*Y ≤ y*) and *p*_*y*_ = *F*_hi_(*y*) *™ F*_lo_(*y*) for the mass at *y*, one observation covers the level *τ* by

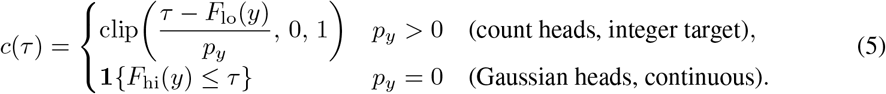

Averaging eq. (5) over the entries of one (slide, gene) pair gives the coverage curve 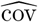, read at 19 levels *τ*_*j*_ equispaced on [0.05, 0.95]. Its residuals 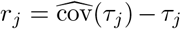 vanish for a calibrated predictive, and two summaries of them are reported,

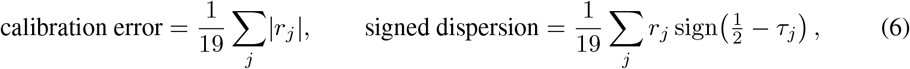

both computed per (slide, gene), then averaged over slides and genes to give one value per held-out donor. The first is the size of the miscalibration, the second its direction, positive when the predictive is too narrow and negative when it is too wide, the sign convention of the signed quantile calibration error [47], to which it is proportional on this symmetric grid. Because a large fraction of entries are zero, at *y* = 0 the coverage contributes only over [0, P(*Y* = 0)], so for the sparser genes a count model’s verdict is largely a statement about how well it predicts the zero mass.

### The marginal baseline

Each family is given a no-image counterpart, which fixes what the choice of distribution already provides before any histology is used. For a held-out donor and a gene, its parameters are estimated on the expression of that gene pooled over the spots of every other donor, on the same target as the trained head of that family, the log target for the Gaussian and the common-depth integer counts for the count heads. The Gaussian baseline takes the pooled mean and variance, the Poisson the pooled mean 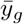, and the negative binomial the pooled mean with the dispersion taken from the pooled variance 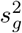 by moments,

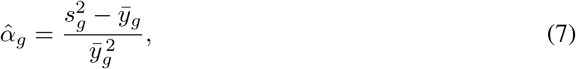

floored at a small positive value. The resulting distribution is used unchanged at every spot of the held-out donor and scored exactly as a trained model, through eq. (5) and the same aggregation over slides and genes.

### Error and uncertainty

The uncertainty of a prediction is the standard deviation of its predictive: *σ* for the Gaussian heads, 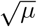 for the Poisson, 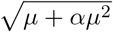 for the negative binomial. Every statistic is a rank correlation, so a variance and its square root give the same value. Two error definitions are used, the absolute error and the absolute log-ratio, the latter defined on positive observations only. An error taken relative to the predicted *spread* is deliberately avoided, since it would make the error a function of the quantity under test.

The absolute log-ratio also requires a positive prediction. Where 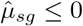 the prediction is replaced by the first percentile of the positive observations of that cohort rather than dropped, since a non-positive prediction is precisely where the model is wrong. A value at the low end of the observed range is chosen so that the substitution does not inflate the error, while still accounting for model incorrectness. It applies to 0.4% to 2.9% of the positive entries depending on the cohort, and only to the Gaussian heads. The count means are the exponentiated output of the MLP and therefore positive. The Gaussian and count families are not scored on the same entries under this error, since the count target rounds the smallest non-zero values to zero, so error levels are read within a family and only the rank correlations are compared across families.

### Interpretation of statistical uncertainty

Each grouping holds 10^4^ to 10^5^ entries, so statistical significance is uninformative here; we take the donor-level *t* interval as the resolution of the measurement. We treat estimates whose confidence intervals include zero as effectively zero and do not interpret differences smaller than the interval half-width. For the controlled correlations of fig. 3, that half-width is 0.005 to 0.031 across the spots of a gene (median 0.012), and 0.008 to 0.077 across the genes of a spot (median 0.043). It stays below 0.018 and 0.055, respectively on Tumor Profiler and NSCLC, and reaches those maxima on Kidney/Lung TLS, the cohort with the fewest donors.

### Controlling for the predicted mean

A group of eq. (2) is retained when it holds at least 60 entries. Groups are averaged inside each held-out donor, and the *K* donor values give the reported mean and its interval. Being a Spearman partial correlation, it removes the monotone part of the dependence on the predicted mean and nothing more, so a residual association surviving it is not attributable to any increasing function of the mean.

### Sparsification

Predictions are withheld in decreasing order of a score at 19 fractions from 0% to 90%, and the mean error of the remainder is recomputed. Removal happens inside a (gene, slide) group, under two protocols: within the group, and within each equal-count bin of the predicted mean inside the group. The latter tests whether the uncertainty tracks the error at a fixed predicted mean. Bins are equal-count quantile bins of the predicted mean, built inside each gene and slide group. The number of bins follows the number of entries of that group: three bins from 60 to 149 entries, five from 150 to 499, and ten from 500 on. Groups holding fewer than 60 entries are excluded. One curve is produced per (gene, slide), normalized by that group’s own mean error, averaged over the slides of a gene weighted by the number of entries, then averaged over genes. Curves are computed twice: on all entries, and on the detected entries alone. On the full panel the error at an undetected entry is the prediction itself, so any score that follows the predicted mean partly sorts by the error.

Four scores are compared: the predicted uncertainty, the true absolute error, which defines that head’s own oracle, and the predicted mean alone in each direction. We report the area under the sparsification error curve and the share of the oracle’s reduction that a score achieves,

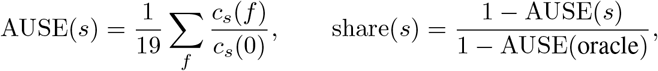

where *c*_*s*_(*f*) is the curve of score *s* at removal fraction *f* . An uninformative ranking gives AUSE 1

### Share of variance decomposition

This is the decomposition reported in section 4.5. For one slide, the retained entries *O* form a spots-by-genes matrix, decomposed on the within-slide ranks *R*_*sg*_. Ranks are used so that the decomposition acts on the same quantities as the Spearman correlation, and ranking within a slide leaves the order of every group unchanged. An additive model *R*_*sg*_ = *m* + *a*_*g*_ + *b*_*s*_ + *r*_*sg*_ is fitted by least squares over those entries, with *m* the mean over *O, a*_*g*_ a gene effect, *b*_*s*_ a spot effect, and *r*_*sg*_ the interaction,

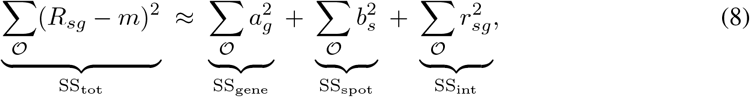

approximate because the detected entries form an unbalanced design. Each term over SS_tot_ gives the reported share, averaged over the slides of a donor and then over donors.

## F Post hoc recalibration

A scale factor on the predicted variance, Var^*′*^ = *c* Var, is fitted on the validation patient of each fold and applied unchanged to the held-out patient. Each gene is cut into ten quantile bins of its own predicted mean, or of its own predicted uncertainty [41]. The bins are computed on the validation patient. The factor minimizes the negative log-likelihood over the bin, in closed form for the Gaussian heads and by a grid scan for the negative binomial.

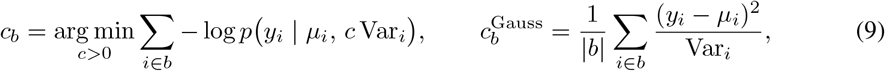

where *b* is a (gene, bin) cell and the second expression is the closed-form minimizer for the Gaussian heads. For the negative binomial the scan is restricted to *c* Var_*i*_ ≥ *µ*_*i*_, since no negative binomial has variance below its mean. Recalibration is applied to the heteroscedastic Gaussian and negative binomial heads. It is not applied to the Poisson head, whose variance is tied to its mean. Calibration error improves by at most 8% and no variant improves any uncertainty-error correlation.

## Footnotes

6 https://github.com/BiomedicalMachineLearning/STimage

## References

[1] Patrik L. Ståhl, Fredrik Salmén, Sanja Vickovic, Anna Lundmark, José Fernández Navarro, Jens Magnusson, Stefania Giacomello, Michaela Asp, Jakub O. Westholm, Mikael Huss, Annelie Mollbrink, Sten Linnarsson, Simone Codeluppi, Åke Borg, Fredrik Pontén, Paul Igor Costea, Pelin Sahlén, Jan Mulder, Olaf Bergmann, Joakim Lundeberg, and Jonas Frisén. Visualization and analysis of gene expression in tissue sections by spatial transcriptomics. Science, 353(6294): 78–82, 2016. doi: 10.1126/science.aaf2403.

[2] Bryan He, Ludvig Bergenstråhle, Linnea Stenbeck, Abubakar Abid, Alma Andersson, Åke Borg, Jonas Maaskola, Joakim Lundeberg, and James Zou. Integrating spatial gene expression and breast tumour morphology via deep learning. Nature Biomedical Engineering, 4(8):827–834, 2020. doi: 10.1038/s41551-020-0578-x.

[3] Yuansong Zeng, Zhuoyi Wei, Weijiang Yu, Rui Yin, Yuchen Yuan, Bingling Li, Zhonghui Tang, Yutong Lu, and Yuedong Yang. Spatial transcriptomics prediction from histology jointly through Transformer and graph neural networks. Briefings in Bioinformatics, 23(5):bbac297, 2022. doi: 10.1093/bib/bbac297.

[4] Kalin Nonchev, Sebastian Dawo, Karina Silina, Holger Moch, Sonali Andani, Tumor Profiler Consortium, Viktor H. Koelzer, and Gunnar Rätsch. DeepSpot: Leveraging spatial context for enhanced spatial transcriptomics prediction from H&E images, 2025. URL https://www.medrxiv.org/content/10.1101/2025.02.09.25321567v2.

[5] Kalin Nonchev, Sebastian Dawo, Karina Silina, Viktor H. Koelzer, and Gunnar Rätsch. DeepSpot-M: a multimodal foundation model for transcriptome-wide virtual spatial transcriptomics from histology, 2026. URL https://www.medrxiv.org/content/10.64898/2026.06.19.26356060v1.

[6] Manuel Tran, Rushin H. Gindra, Philipp Putze, Kang Senbai, Giovanni Palla, Tina Kos, Chiara Falcomatà, Chen Wang, Ruifeng (Ray) Guo, Melanie Boxberg, Luc M. Berclaz, Lars H. Lindner, Linda Bergmayr, Thomas Knösel, Philipp Jurmeister, Frederick Klauschen, Krisztian Homicsko, Raphael Gottardo, Markus Eckstein, Christian Matek, Andreas Mock, Fabian J. Theis, Dieter Saur, and Tingying Peng. Pan-cancer virtual spatial transcriptomics from routine histology with Phoenix, 2026. URL https://www.biorxiv.org/content/10.64898/2026.04.25.720812v1.

[7] Daiwei Zhang, Amelia Schroeder, Hanying Yan, Haochen Yang, Jian Hu, Michelle Y. Y. Lee, Kyung S. Cho, Katalin Susztak, George X. Xu, Michael D. Feldman, et al. Inferring superresolution tissue architecture by integrating spatial transcriptomics with histology. Nature Biotechnology, 42(9):1372–1377, 2024. doi: 10.1038/s41587-023-02019-9.

[8] Kalin Nonchev, Glib Manaiev, Viktor H. Koelzer, and Gunnar Rätsch. DeepSpot2Cell: predicting virtual single-cell spatial transcriptomics from H&E images using spot-level supervision, 2025.

[9] Yuanhao Jiang, Jacky Xie, Xiao Tan, Nan Ye, and Quan Nguyen. Generalization of deep learning models for predicting spatial gene expression profiles using histology images: A breast cancer case study, 2023. URL https://www.biorxiv.org/content/10.1101/2023.09.20.558624v1.

[10] Christian Leibig, Vaneeda Allken, Murat Seçkin Ayhan, Philipp Berens, and Siegfried Wahl. Leveraging uncertainty information from deep neural networks for disease detection. Scientific Reports, 7:17816, 2017. doi: 10.1038/s41598-017-17876-z.

[11] James M. Dolezal, Andrew Srisuwananukorn, Dmitry Karpeyev, Siddhi Ramesh, Sara Kochanny, Brittany Cody, Aaron S. Mansfield, Sagar Rakshit, Radhika Bansal, Melanie C. Bois, Aaron O. Bungum, Jefree J. Schulte, Everett E. Vokes, Marina Chiara Garassino, Aliya N. Husain, and Alexander T. Pearson. Uncertainty-informed deep learning models enable high-confidence predictions for digital histopathology. Nature Communications, 13(1):6572, 2022. doi: 10.1038/s41467-022-34025-x.

[12] Max-Heinrich Laves, Sontje Ihler, Jacob F. Fast, Lüder A. Kahrs, and Tobias Ortmaier. Well-calibrated regression uncertainty in medical imaging with deep learning. In Proceedings of the Third Conference on Medical Imaging with Deep Learning (MIDL), volume 121 of Proceedings of Machine Learning Research, pages 393–412. PMLR, 2020.

[13] Lior Hirschfeld, Kyle Swanson, Kevin Yang, Regina Barzilay, and Connor W. Coley. Uncertainty quantification using neural networks for molecular property prediction. Journal of Chemical Information and Modeling, 60(8):3770–3780, 2020. doi: 10.1021/acs.jcim.0c00502.

[14] Xiao Tan, Onkar Mulay, Jacky Xie, Samual MacDonald, Taehyun Kim, Chenhao Zhou, Zherui Xiong, Samuel X. Tan, Nan Ye, Amy McCart Reed, Kiarash Khosrotehrani, Fred Roosta, Maciej Trzaskowski, Peter T. Simpson, and Quan Nguyen. Robust and interpretable prediction of gene markers and cell types from spatial transcriptomics data. Nature Communications, 17:1781, 2026. doi: 10.1038/s41467-026-68487-0.

[15] Thanosan Prathifkumar, Nehan Mohamed, Anvith Kakkera, Zarif Azher, Xiaoying Liu, Fred Kolling, Laurent Perreard, Scott Palisoul, Louis J. Vaickus, and Joshua Jay Levy. Quantifying uncertainty in virtual spatial transcriptomics using Bayesian neural networks. Cancer Research, 86(7 Suppl.):4189, 2026. doi: 10.1158/1538-7445.AM2026-4189. Abstract nr 4189, AACR Annual Meeting 2026.

[16] Michael I. Love, Wolfgang Huber, and Simon Anders. Moderated estimation of fold change and dispersion for RNA-seq data with DESeq2. Genome Biology, 15(12):550, 2014. doi: 10.1186/s13059-014-0550-8.

[17] Mark D. Robinson, Davis J. McCarthy, and Gordon K. Smyth. edgeR: a Bioconductor package for differential expression analysis of digital gene expression data. Bioinformatics, 26(1): 139–140, 2010. doi: 10.1093/bioinformatics/btp616.

[18] Romain Lopez, Jeffrey Regier, Michael B. Cole, Michael I. Jordan, and Nir Yosef. Deep generative modeling for single-cell transcriptomics. Nature Methods, 15(12):1053–1058, 2018. doi: 10.1038/s41592-018-0229-2.

[19] Tim Stuart, Andrew Butler, Paul Hoffman, Christoph Hafemeister, Efthymia Papalexi, III Mauck, William M., Yuhan Hao, Marlon Stoeckius, Peter Smibert, and Rahul Satija. Comprehensive integration of single-cell data. Cell, 177(7):1888–1902.e21, 2019. doi: 10.1016/j.cell.2019.05.031.

[20] Srikar Yellapragada, Alexandros Graikos, Zilinghan Li, Kostas Triaridis, Varun Belagali, Saarthak Kapse, Tarak Nath Nandi, Ravi K. Madduri, Prateek Prasanna, Tahsin Kurc, Rajarsi R. Gupta, Joel Saltz, and Dimitris Samaras. PixCell: A generative foundation model for digital histopathology images, 2025. URL https://arxiv.org/abs/2506.05127.

[21] Pantelis R. Vlachas, Kalin Nonchev, Viktor H. Koelzer, and Gunnar Rätsch. Transcriptomics-conditioned virtual tissue synthesis via diffusion transformers. bioRxiv, 2026.

[22] Moritz Schaefer, Kalin Nonchev, Animesh Awasthi, Jake Burton, Viktor H. Koelzer, Gunnar Rätsch, and Christoph Bock. Molecularly informed analysis of histopathology images using natural language. bioRxiv, 2025.

[23] Moritz Schaefer, Zoe Piran, Nils Philipp Walter, Animesh Awasthi, Christoph Bock, Jure Leskovec, and Zinaida Good. Transitive representation learning enhances histopathology annotation. In Forty-third International Conference on Machine Learning, 2026.

[24] Alex Kendall and Yarin Gal. What uncertainties do we need in Bayesian deep learning for computer vision? In Advances in Neural Information Processing Systems, volume 30, pages 5574–5584, 2017.

[25] Balaji Lakshminarayanan, Alexander Pritzel, and Charles Blundell. Simple and scalable predictive uncertainty estimation using deep ensembles. In Advances in Neural Information Processing Systems, volume 30, pages 6402–6413, 2017.

[26] Yarin Gal and Zoubin Ghahramani. Dropout as a Bayesian approximation: Representing model uncertainty in deep learning. In Proceedings of the 33rd International Conference on Machine Learning (ICML), volume 48 of Proceedings of Machine Learning Research, pages 1050–1059. PMLR, 2016. URL https://proceedings.mlr.press/v48/gal16.html.

[27] Erik Daxberger, Agustinus Kristiadi, Alexander Immer, Runa Eschenhagen, Matthias Bauer, and Philipp Hennig. Laplace redux – effortless Bayesian deep learning. In Advances in Neural Information Processing Systems, volume 34, pages 20089–20103, 2021.

[28] Volodymyr Kuleshov, Nathan Fenner, and Stefano Ermon. Accurate uncertainties for deep learning using calibrated regression. In Proceedings of the 35th International Conference on Machine Learning (ICML), volume 80 of Proceedings of Machine Learning Research, pages 2796–2804. PMLR, 2018. URL https://proceedings.mlr.press/v80/kuleshov18a.html.

[29] Tilmann Gneiting, Fadoua Balabdaoui, and Adrian E. Raftery. Probabilistic forecasts, calibration and sharpness. Journal of the Royal Statistical Society: Series B (Statistical Methodology), 69 (2):243–268, 2007. doi: 10.1111/j.1467-9868.2007.00587.x.

[30] Eric D. Sun, Rong Ma, Paloma Navarro Negredo, Anne Brunet, and James Zou. TISSUE: uncertainty-calibrated prediction of single-cell spatial transcriptomics improves downstream analyses. Nature Methods, 21(3):444–454, 2024. doi: 10.1038/s41592-024-02184-y.

[31] Kaitian Jin, Zihao Chen, Xiaokang Yu, Musu Yuan, Amelia Schroeder, Bernhard Dumoulin, Yunhe Liu, Linghua Wang, Jeong Hwan Park, Tae Hyun Hwang, Katalin Susztak, Zhimei Ren, Nancy R. Zhang, and Mingyao Li. Multiscale confidence quantification for virtual spatial transcriptomics with UTOPIA, 2026. URL https://www.biorxiv.org/content/10.64898/2026.03.01.708850v1.

[32] Mikhail Karasikov, Joost van Doorn, Nicolas Känzig, Melis Erdal Cesur, Hugo Mark Horlings, Robert Berke, Fei Tang, and Sebastian Otálora. Training state-of-the-art pathology foundation models with orders of magnitude less data. In Medical Image Computing and Computer Assisted Intervention – MICCAI 2025, volume 15967 of Lecture Notes in Computer Science, pages 573–583. Springer, 2025. doi: 10.1007/978-3-032-04984-1_55.

[33] Claudia Czado, Tilmann Gneiting, and Leonhard Held. Predictive model assessment for count data. Biometrics, 65(4):1254–1261, 2009. doi: 10.1111/j.1541-0420.2009.01191.x.

[34] Maximilian Seitzer, Arash Tavakoli, Dimitrije Antić, and Georg Martius. On the pitfalls of heteroscedastic uncertainty estimation with probabilistic neural networks. In International Conference on Learning Representations (ICLR), 2022.URL https://openreview.net/forum?id=aPOpXlnV1T.

[35] Lydia A. Schoenpflug, Nikki van den Berg, Sonali Andani, Nanda Horeweg, Jurriaan Barkey Wolf, Tjalling Bosse, Viktor H. Koelzer, and Maxime W. Lafarge. A protocol for evaluating robustness to H&E staining variation in computational pathology models. Journal of Pathology Informatics, page 100702, 2026.

[36] Linnea Stenbeck, Fanny Taborsak-Lines, and Stefania Giacomello. Enabling automated and reproducible spatially resolved transcriptomics at scale. Heliyon, 8(6):e09651, 2022. doi: 10.1016/j.heliyon.2022.e09651.

[37] Mei R. M. Du, Changqing Wang, Charity W. Law, Daniela Amann-Zalcenstein, Casey J. A. Anttila, Ling Ling, Peter F. Hickey, Callum J. Sargeant, Yunshun Chen, Lisa J. Ioannidis, Pradeep Rajasekhar, Raymond K. H. Yip, Kelly L. Rogers, Diana S. Hansen, Rory Bowden, and Matthew E. Ritchie. Benchmarking spatial transcriptomics technologies with the multi-sample SpatialBenchVisium dataset. Genome Biology, 26:77, 2025. doi: 10.1186/s13059-025-03543-4.

[38] Jasmine T. Plummer, Felipe Segato Dezem, David P. Cook, Jiwoon Park, Luke Zhang, Yutian Liu, Maycon Marção, Hannah DuBose, Arjumand Wani, Kellie Wise, Michael Roach, Kate Harvey, Taopeng Wang, Kirk B. Jensen, Natalia Morosini, Roberto De Gregorio, Alicia Alonso, Shauna Lee Houlihan, Robert E. Schwartz, Erika Hissong, Catherine Snopkowski, Jeffrey L. Wrana, Natalie Ryan, Lisa M. Butler, George Church, Alexander Swarbrick, Christopher E. Mason, and Luciano G. Martelotto. Standardized metrics for assessment and reproducibility of imaging-based spatial transcriptomics datasets. Nature Biotechnology, 44:1213–1225, 2026. doi: 10.1038/s41587-025-02811-9.

[39] Anastasios N. Angelopoulos, Amit Pal Kohli, Stephen Bates, Michael I. Jordan, Jitendra Malik, Thayer Alshaabi, Srigokul Upadhyayula, and Yaniv Romano. Image-to-image regression with distribution-free uncertainty quantification and applications in imaging. In Proceedings of the 39th International Conference on Machine Learning (ICML), volume 162 of Proceedings of Machine Learning Research, pages 717–730. PMLR, 2022. URL https://proceedings.mlr.press/v162/angelopoulos22a.html.

[40] Hao Song, Tom Diethe, Meelis Kull, and Peter Flach. Distribution calibration for regression. In Proceedings of the 36th International Conference on Machine Learning (ICML), volume 97 of Proceedings of Machine Learning Research, pages 5897–5906. PMLR, 2019. URL https://proceedings.mlr.press/v97/song19a.html.

[41] Pascal Pernot. Can bin-wise scaling improve consistency and adaptivity of prediction uncertainty for machine learning regression?, 2023. URL https://arxiv.org/abs/2310.11978.

[42] Nicola Miglino, Nora C Toussaint, Alexander Ring, Ximena Bonilla, Marina Tusup, Benedict Gosztonyi, Tarun Mehra, Gabriele Gut, Francis Jacob, Stephane Chevrier, et al. Feasibility of multiomics tumor profiling for guiding treatment of melanoma. Nature Medicine, 31(7): 2430–2441, 2025.

[43] Kalin Nonchev, Sonali Andani, Joanna Ficek-Pascual, Marta Nowak, Bettina Sobottka, Tumor Profiler Consortium, Viktor H. Koelzer, and Gunnar Rätsch. Representation learning for multi-modal spatially resolved transcriptomics data. Bioinformatics, 2026. doi: 10.1093/bioinformatics/btag316.

[44] Sebastian Dawo, Kalin Nonchev, and Karina Silina. 10x Visium spatial transcriptomics dataset: Kidney (3) and lung (5) cancer with tertiary lymphoid structures, 2025. URL 10.5281/zenodo.14620362. Dataset.

[45] Marco De Zuani, Haoliang Xue, Jun Sung Park, Stefan C. Dentro, Zaira Seferbekova, Julien Tessier, Sandra Curras-Alonso, Angela Hadjipanayis, Emmanouil I. Athanasiadis, Moritz Gerstung, Omer Bayraktar, and Ana Cvejic. Single-cell and spatial transcriptomics analysis of non-small cell lung cancer. Nature Communications, 15(1):4388, 2024. doi: 10.1038/s41467-024-48700-8.

[46] Yaniv Ovadia, Emily Fertig, Jie Ren, Zachary Nado, D. Sculley, Sebastian Nowozin, Joshua V. Dillon, Balaji Lakshminarayanan, and Jasper Snoek. Can you trust your model’s uncertainty? Evaluating predictive uncertainty under dataset shift. In Advances in Neural Information Processing Systems, volume 32, pages 13969–13980, 2019.

[47] Florian Seligmann, Philipp Becker, Michael Volpp, and Gerhard Neumann. Beyond deep ensembles: A large-scale evaluation of Bayesian deep learning under distribution shift. In Advances in Neural Information Processing Systems, volume 36, 2023. URL https://proceedings.neurips.cc/paper_files/paper/2023/hash/5d97b7e62022c859347397f6c1e8d0f9-Abstract-Conference.html.

